# PIEZO2 in somatosensory neurons controls gastrointestinal transit

**DOI:** 10.1101/2022.11.27.518109

**Authors:** M. Rocio Servin-Vences, Ruby M. Lam, Alize Koolen, Yu Wang, Dimah N. Saade, Meaghan Loud, Halil Kacmaz, Arthur Beyder, Kara L. Marshall, Carsten G. Bönnemann, Alexander T. Chesler, Ardem Patapoutian

**Affiliations:** Department of Neuroscience, Dorris Neuroscience Center, Scripps Research, San Diego, California, USA; Howard Hughes Medical Institute, Chevy Chase, USA; National Institute of Neurological Disorders and Stroke, National Institutes of Health, Bethesda, Maryland, USA; Division of Gastroenterology and Hepatology, Enteric Neuroscience Program (ENSP), Mayo Clinic, Rochester, Minnesota, USA; Department of Physiology and Biomedical Engineering, Mayo Clinic, Rochester, Minnesota, USA; Department of Neuroscience, Baylor College of Medicine, Jan and Dan Duncan Neurological Research Institute, Houston, Texas, USA; National Center for Complementary and Integrative Health, National Institutes of Health, Bethesda, Maryland, USA

## Abstract

The gastrointestinal tract is in a state of constant motion. These movements are tightly regulated by the presence of food and help digestion by mechanically breaking down and propelling gut content. Mechanical sensing in the gut is thought to be essential for regulating motility; however, the identity of the neuronal populations, the molecules involved, and the functional consequences of this sensation are unknown. Here, we show that humans lacking PIEZO2 exhibit impaired bowel sensation and motility. Piezo2 in mouse dorsal root but not nodose ganglia is required to sense gut content, and this activity slows down food transit rates in the stomach, small intestine, and colon. Indeed, Piezo2 is directly required to detect colon distension *in vivo*. Our study unveils the mechanosensory mechanisms that regulate the transit of luminal contents throughout the gut, which is a critical process to ensure proper digestion, nutrient absorption, and waste removal.

**Highlights:**

- Individuals with PIEZO2 syndrome present impaired bowel sensation and gastrointestinal dysfunction.
- PIEZO2 in DRG neurons plays an important role in regulating gut motility.
- Lack of PIEZO2 from sensory neurons accelerates gastric emptying and intestinal transit.
- DRG neurons detect colon distension via PIEZO2.

## Introduction

Neural mechanisms regulate key functions of the gastrointestinal (GI) tract, including motility which is necessary to break down the ingested food, to absorb its components and to eliminate waste ^1^. After swallowing, food moves in an orderly way through specialized compartments, each with distinct functions. Thus, the propulsion of gut contents is tightly regulated. Throughout the GI tract, mechanical mixing serves as a key process to enhance efficiency of chyme breakdown and keep ingested contents moving ^2^. Well defined efferent motor programs mediate gut motility through stereotyped movements (e.g peristalsis, segmentation and ‘migrating motor complexes’^1–3^) that are initiated and controlled by complex neural inputs that respond to chemical and mechanical stimuli ^4–6^. However, little is known about the molecular mechanisms that coordinate and initiate motility along the GI tract, including the molecular identity of mechanosensors within the gut, as well as the key sensory neurons that modulate motility along the GI tract.

There are three major afferent neural pathways in the gut. The Enteric Nervous System (ENS) is intrinsic to the gut and functions to initiate local motility reflexes ^5,7^. Vagal neurons from the Nodose ganglion and somatosensory neurons from dorsal root ganglia (DRGs) are extrinsic to the GI tract, yet both richly innervate the gut ^8–10^. It is generally accepted that nodose neurons play key roles in mediating homeostatic gut-brain signaling ^11–14^ whereas DRG neurons are critically important for sensing gut inflammation and evoking pain ^15,16^. A conserved feature of all three gut afferent systems is that they contain neurons that detect and respond to chemical and mechanical stimuli ^8,17–20^. However, many of these studies are performed *in situ*, at the whole-ganglion level, and do not distinguish the specific role and outcomes of mechanosensation versus chemosensation. Furthermore, far less is known about the molecular mechanisms that control the transit of ingested contents along the GI tract *in vivo*.

PIEZO2 is a mechanically gated ion channel that is the receptor for gentle touch and proprioception in mice and humans ^21–23^. More recently, work of our group and others have shown that PIEZO2 also has critical functions in interoception, including sensing lung inflation ^24^ and sensing bladder filling ^25^. Thus, expression of PIEZO2 is one of the strongest predictors of a cell having a critical mechanosensory role. Notably, this molecule is expressed in all three gut-innervating neural systems-the enteric ^18,19^, vagal ^8,24,26,27^ and somatosensory systems ^23,28,29^-yet its function in any of these systems is unknown. Here, we collected clinical data from a group of *PIEZO2*-deficient individuals and used genetic mouse models to interrogate the role of Piezo2 in gut transit.

## Results

### Gastrointestinal dysfunction in individuals deficient in *PIEZO2*

To better understand the role of PIEZO2 in human GI function, we assessed the GI health and medical history of human subjects carrying *PIEZO2* loss-of-function variants (n = 8; ages 9 to 42). We additionally used PROMIS (Patient Reported Outcomes Measurement Information System) questionnaires, a clinical tool developed by the National Institute of Health to capture and evaluate general GI symptoms ^30,31^. These GI questionnaires are widely used as patient-reported health information and capture answers from the previous seven days to the survey. The responses obtained from the *PIEZO2*-deficienct individuals were compared with the 1,177 control answers from general-population volunteers ^30,31^ (Figure 1). We observed GI dysfunctions that changed with age: *PIEZO2*-deficient children frequently reported lumpy stools, teenagers had lumpy and watery stools, and older adults tended to have watery stools (Figure 1; 7 subjects answered the survey). Additionally, *PIEZO2*-deficient children reported needing constant strain during bowel movements, whereas older individuals had a sudden urgency to evacuate their bowels. Eight individuals reported childhood constipation that improved or disappeared with age, and the oldest adult (42 years old) reported having recurrent diarrhea that was improved with dietary changes. Remarkably, six *PIEZO2*-deficient subjects reported difficulties in sensing bowel movements, instead, they determined successful stool passage by relying on sound, smell, and/or vision. Three individuals follow a specific daily bowel regimen to cope with their lack of bowel movement sensation, while three other individuals reported soiling accidents. Additionally, five patients reported taking medication to aid with GI distress. Although access to *PIEZO2*-deficient individuals is rare, the captured information allowed us to formulate hypothesis regarding the role of PIEZO2 in GI function. Thus, these findings indicate that *PIEZO2*-deficient individuals have impaired sensation in bowel function that affects their quality of life and suggest that the mechanosensitive channel PIEZO2 plays a crucial role in human GI physiology and pathophysiology.

**Figure 1.**
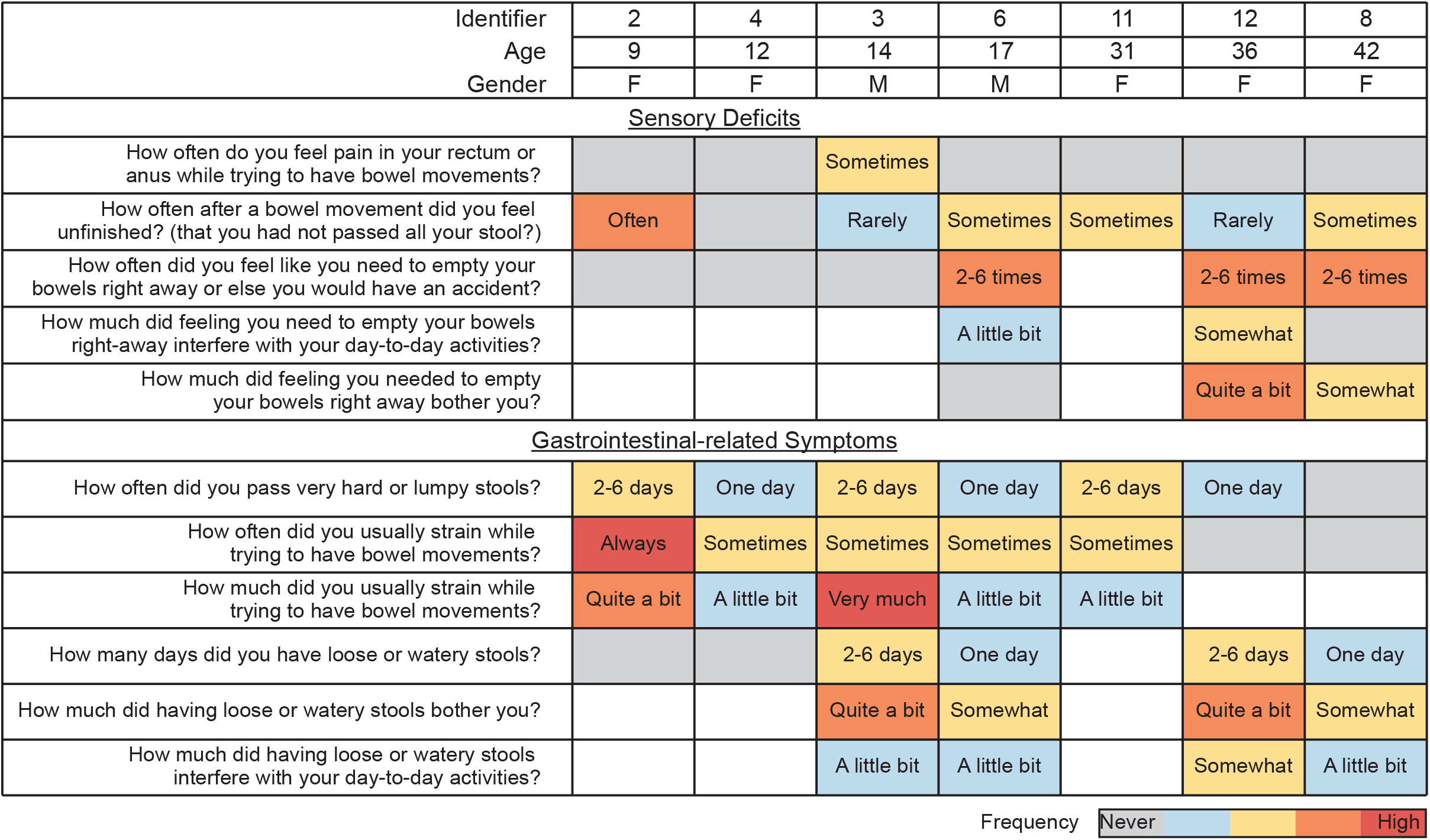
Gastrointestinal dysfunction in individuals deficient in *PIEZO2*. Summary of responses obtained from *PIEZO2*-individuals to GI-PROMIS questionnaires. Data indicates the Patient identifier, age at which the questionnaires were answered and gender. Data is organized by ascending age and symptoms are divided between sensory and GI, which span constipation and diarrhea. Each question assessed symptoms from the seven days prior to the survey. Unless otherwise noted, the color code indicates the following: grey: never -no pathology-the average answer of the healthy control individuals; blue: rarely; yellow: sometimes; orange: often, red: always. Therefore, every color except for gray indicates a deviation from the average. Blank indicates unanswered question. Individual identifier corresponds to those published in our previous urinary function study ^25^.

### Piezo2 in sensory neurons is required for gastrointestinal function in mice

Intrinsic and extrinsic neuronal innervation of the gut are essential for normal GI motility. Vagotomies commonly result in delayed gastric emptying ^3,32^, lack of ENS results in Hirschsprung’s disease that causes the inability to pass stool through the colon ^33^, and spinal cord injuries often lead to fecal incontinence and constipation ^34^. In order to establish the role of neuronal Piezo2 in GI physiology, we used transgenic mouse models to ablate Piezo2 from peripheral sensory neurons. We used the *Scn10a*^*Cre*^ driver line (*SNS*^*Cre* 35^) which expresses Cre recombinase under the regulatory elements of the *Scn10a* gene (voltage gated sodium channel Na_v_1.8). First, we established the extent of recombination in the three sources of gut innervation: enteric, DRG and vagal. Previous reports had shown that the *SNS*^*Cre*^ driver recombines in about 80% of neurons from the vagal and DRG ^35–37^, but not in other cell types such as enterochromaffin cells ^37^. However, there is little information about its efficiency in enteric neurons along the GI tract. To validate the *SNS*^*Cre*^ dependent recombination in the ENS, we crossed *SNS*^*Cre+/-*^ mice to *Ai9*^*/fl/fl*^ mice and detected partial signal along the GI tract (Figure S1-A). To verify that there is minimal overlap between *Piezo2* expression in *Scn10* enteric neurons, we mined a single-cell transcriptomic data set ^38^ and observed no coexpression of *Piezo2* and *Scn10* transcripts (Figure S1-B). Given these results, we do not anticipate that phenotypes in this line are a result of *Piezo2* expression in the ENS.

Next, we studied the effects of Piezo2 deletion in GI function, by evaluating the GI transit time, evacuation frequency, and stool water content of *SNS*^*Cre*^ crossed *Piezo2*^*fl/fl*^ mice (Figure 2A). To measure whole GI transit, we gavaged mice with carmine red, a non-absorbable red dye with no nutritional value. We then recorded the time for the first appearance of colored feces. We observed a robust transit time acceleration in the conditional knockout (*SNS*^*Cre+/-*^*;Piezo2*^*fl/fl*^, referred to here as *Piezo2*^*SNS*^) mice, compared to the wild-type (*SNS*^*Cre-/-*^*;Piezo2*^*fl/fl*^, *Piezo2*^*WT*^) littermate controls (Figure 2B). Notably, Piezo2 deletion did not affect small intestine and colon length (Figure S1C, D). Moreover, the *Piezo2*^*SNS*^ mice expelled a greater number of stools during one hour of sample collection and presented a significant increase in stool water content in comparison with the *Piezo2*^*WT*^ littermates (Figure 2C-E). In agreement with the higher amount of water content, the dried-stool weight from the *Piezo2*^*SNS*^ mice was significantly smaller in comparison with the *Piezo2*^*WT*^ controls (Figure 2C, F), suggesting that the accelerated transit did not allow time for adequate water absorption. These results indicate that the *Piezo2*^*SNS*^ mice have accelerated GI transit resulting in shorter transit time and a diarrhea-like phenotype.

**Figure 2.**
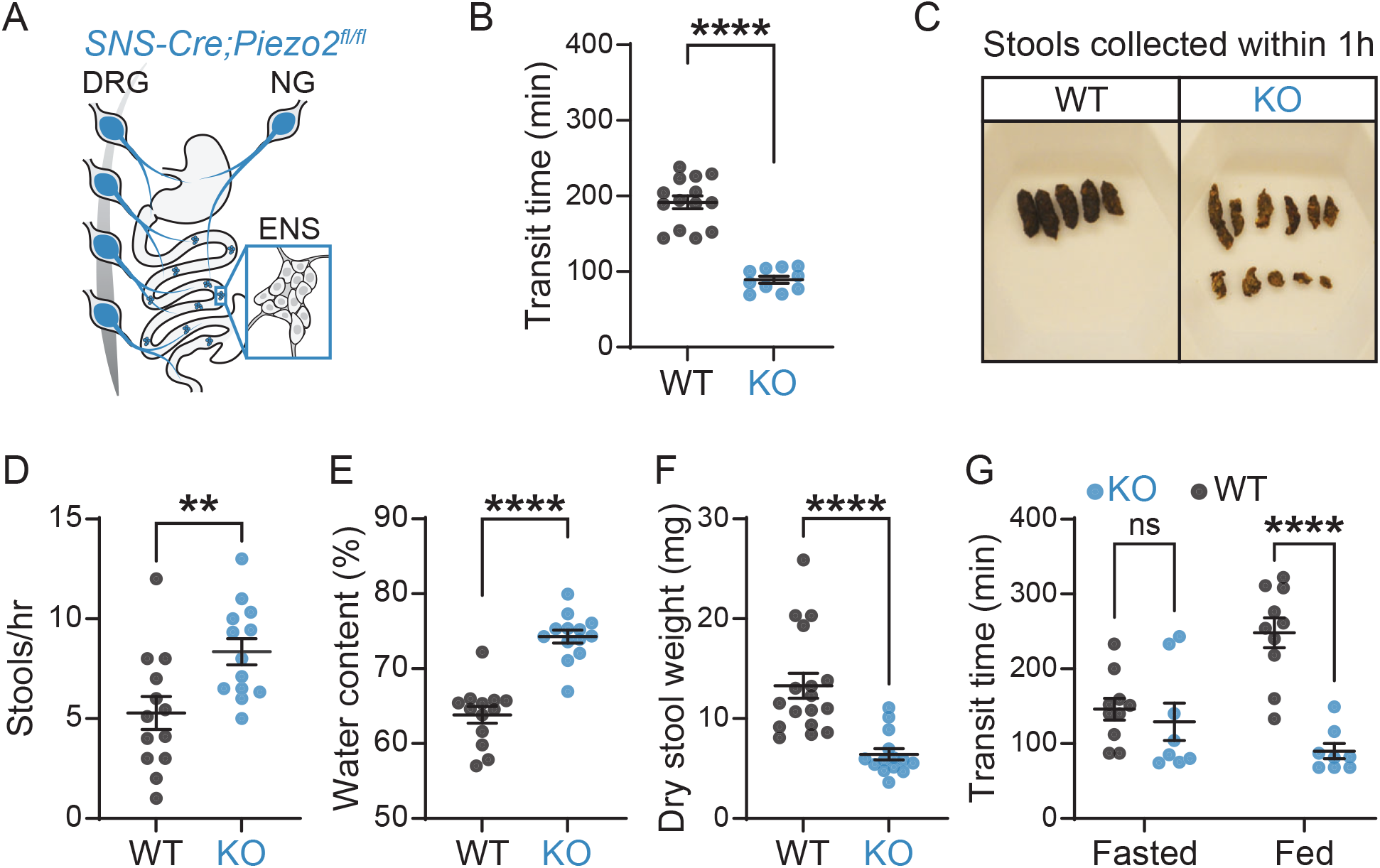
Piezo2 in sensory neurons is required for gastrointestinal function in mice. A) Illustration of the *SNS-Cre;Piezo2*-targeting coverage of extrinsic neurons that innervate the GI tract. Blue designates *Piezo2* deletion in nodose and DRG neurons, but not ENS. B) Total GI transit time measured after gavaging carmine red dye into *SNS-Cre*^*-/-*^*;Piezo2*^*fl/fl*^ (WT; n=14) and *SNS-Cre*^*+/-*^*;Piezo2*^*fl/fl*^ (KO; n=10) mice (unpaired two-tailed *t*-test: \*\*\*\**P*<0.0001, t(22)=9.301). C) Representative images of dried stools collected during one hour from *SNS-Cre*^*-/-*^*;Piezo2*^*fl/fl*^ (WT) and *SNS-Cre*^*+/-*^*;Piezo2*^*fl/fl*^ (KO) mice. D) Number of stools expelled during 1 hr of collection from *SNS-Cre*^*-/-*^*;Piezo2*^*fl/fl*^ (WT; n=13) and *SNS-Cre*^*+/-*^*;Piezo2*^*fl/fl*^ (KO; n=13) mice (unpaired two-tailed *t*-test: \*\**P*=0.0076, t(24)=2.916). E) Water content present in the stool samples from panel (**D**) as a percent of the total composition (unpaired two-tailed *t*-test: \*\*\*\**P*<0.0001, t(24)=7.418). F) All the stool samples collected in panel (**D**) were dried, individually weighted and averaged per mouse: *SNS-Cre*^*-/-*^*;Piezo2*^*fl/fl*^ (WT; n=17) and *SNS-Cre*^*+/-*^*;Piezo2*^*fl/fl*^ (KO; n=14) (Mann-Whitney test: \*\*\*\**P*<0.0001 two-tailed, U=16). G) Total GI transit time measured after gavaging carmine red dye in mice fasted for 12 hours or with *ad libitum* food access. *SNS-Cre*^*-/-*^*;Piezo2*^*fl/fl*^ (WT; n=10) and *SNS-Cre*^*+/-*^*;Piezo2*^*fl/fl*^ (KO; n=8) mice (two-way ANOVA: \*\*\*\**P*_genotype_=0.0009, *F*(1,16)=16.32; Sidak’s *P*_adjusted_: *P*_Fasted_=0.7697; \*\*\*\**P*_Fed_<0.0001).

Finally, to investigate whether the presence of intestinal contents is important to modulate the quickening of the GI transit, we compared gut transit between mice fasted for 12 hours and mice fed *ad libitum*, which already have food contents along the GI tract. Interestingly, we did not observe any transit difference in fasted *Piezo2*^*SNS*^ and *Piezo2*^*WT*^ mice (Figure 2G). Importantly, these results suggest that the mechanical signals exerted by the intestinal contents are directly or indirectly sensed by Piezo2 to modulate GI transit *in vivo*. Moreover, these results indicate that Piezo2 in sensory neurons slow gut transit during digestion periods. Thus, subsequent experiments were performed in *ad libitum* condition.

### Piezo2 in somatosensory neurons is required for gastrointestinal transit in mice

Piezo2 is expressed in cells that influence GI motility, including extrinsic neurons of spinal and vagal origin that innervate the gut ^8,28^, and in enterochromaffin cells of the small intestine and colon ^39,40^. We undertook a targeted approach utilizing genetic and viral methods to identify the specific contributions of Piezo2 -dependent mechanotransductioin in gut transit. We used *Phox2b*^*Cre*^ and *Vil1*^*Cre*^ to target nodose neurons and gut epithelial cells respectively, as well as *Hoxb8*^*Cre*^ to target both caudal DRGs and gut epithelial cells, and finally we used viral intrathecal injections of Cre recombinase to target only DRGs neurons.

Previous studies have demonstrated the importance of nodose innervation in the GI function ^9,10,14,41^. To investigate if vagal sensory neurons could be controlling the faster GI transit seen in the *Piezo2*^*SNS*^ mice, we employed the *Phox2b*^*Cre*^ driver line. As *Phox2b* transcript is widely detected in enteric neurons ^18,19^, we crossed the *Ai9*^*fl/fl*^ reporter mice to the *Phox2b*^*Cre*^ driver to validate the recombination in the ENS. We observed sparse labeling through the gut (Figure S2A), suggesting that in our hands and for our purpose, this *Phox2b*^*Cre*^ driver mainly targets the nodose ganglia. To evaluate the mechanosensory role of vagal innervation in GI transit time, evacuation frequency, and stool water content, we deleted Piezo2 from nodose neurons by crossing of *Phox2b*^*Cre*^ to *Piezo2*^*fl/fl*^ mice. Surprisingly, we found that *Phox2b*^*Cre+/*^*;Piezo2*^*fl/fl*^ (*Piezo2*^*Phox2b*^) mice did not show any difference in transit time and defecation frequency in comparison to their wild-type littermate controls (*Phox2b*^*Cre-/-*^ *;Piezo2*^*fl/fl*^, *Piezo2*^*WT*^) (Figure 3A). Consistent with this finding, the water content and dry-stool weight from *Piezo2*^*Phox2b*^ mice were similar to the *Piezo2*^*WT*^ littermates (Figure S2-D). This indicates that loss of Piezo2 in vagal sensory neurons is insufficient to cause the accelerated GI transit observed in the *Piezo2*^*SNS*^ model.

**Figure 3.**
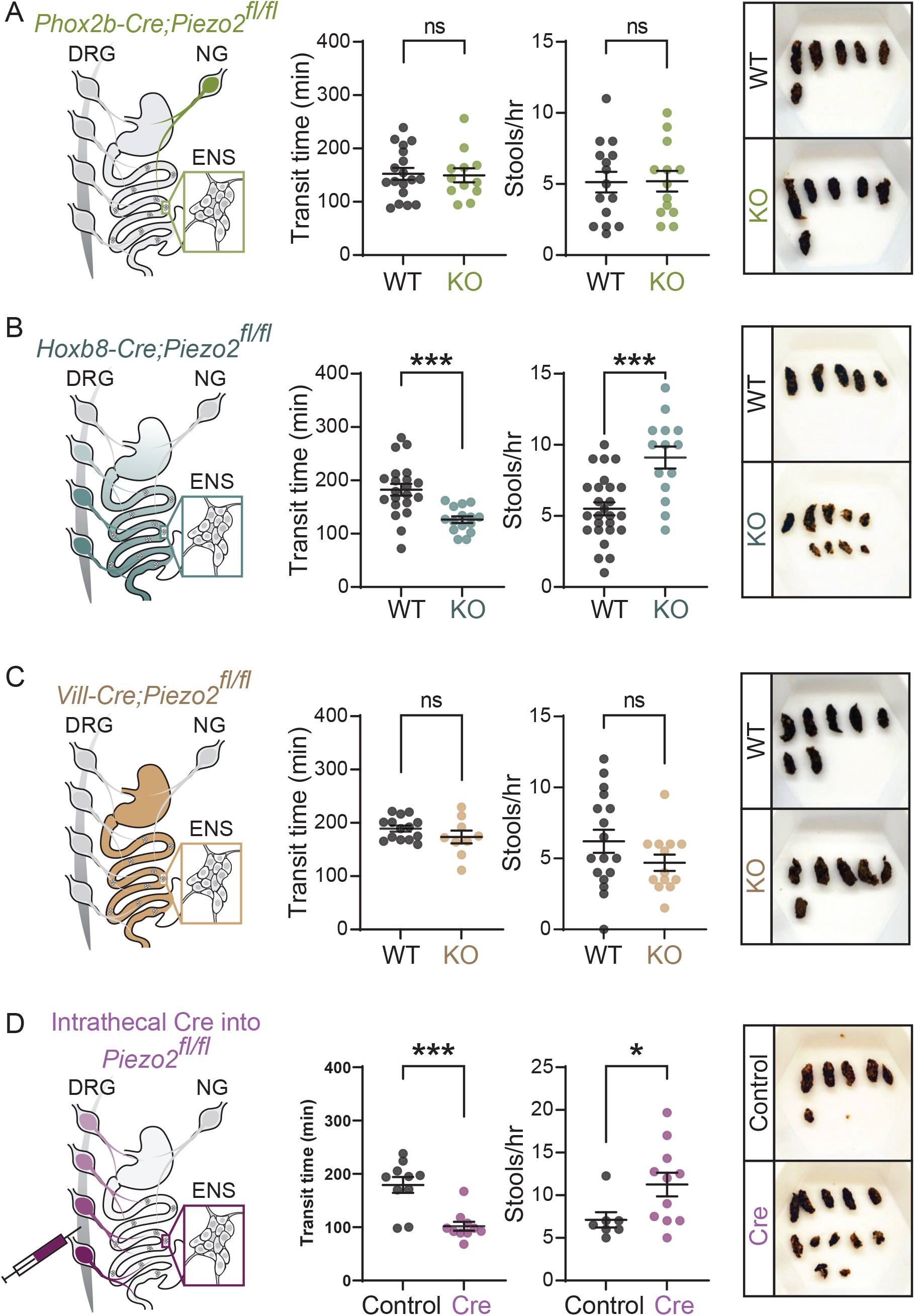
Piezo2 in DRG neurons is required for gastrointestinal transit in mice. A) Illustration of the *Phox2b-Cre;Piezo2* targeting coverage in neurons innervating the GI tract, green designates *Piezo2* deletion in nodose, but not in DRG and enteric neurons (left panel). Total GI transit time after gavaging carmine red into *Phox2b-Cre*^*-/-*^*;Piezo2*^*fl/fl*^ (WT; n=18) and *Phox2b-Cre*^*+/-*^*;Piezo2*^*fl/fl*^ (KO; n=12) mice (unpaired two-tailed *t*-test: *P*=0.8735, t(28)=0.1607; not statistically significant) (middle left panel). Number of stools expelled during 1 hr of collection from *Phox2b-Cre*^*-/-*^*;Piezo2*^*fl/fl*^ (WT; n=15) and *Phox2b-Cre*^*+/-*^*;Piezo2*^*fl/fl*^ (KO; n=13) mice (unpaired two-tailed *t*-test: *P*=0.9548, t(26)=0.05727; not statistically significant) (middle right panel). Representative images of dried stools collected during one hour from *Phox2b-Cre*^*-/-*^*;Piezo2*^*fl/fl*^ (WT) and *Phox2b-Cre*^*+/-*^*;Piezo2*^*fl/fl*^ (KO) mice (right panel). B) Illustration of the *Hoxb8-Cre;Piezo2* targeting coverage in the gastrointestinal epithelium and neurons innervating the gastrointestinal tract, teal color designates *Piezo2* deletion in DRG neurons and enterochromaffin cells of intestinal epithelia, but not in enteric and nodose neurons (left panel). Total gastrointestinal transit time after gavaging carmine red into *Hoxb8-Cre*^*-/-*^*;Piezo2*^*fl/fl*^ (WT; n=21) and *Hoxb8-Cre*^*+/-*^*;Piezo2*^*fl/fl*^ (KO; n=15) mice (unpaired two-tailed *t*-test: \*\*\**P*=0.0003, t(34)=4.004) (middle left panel). Number of stools expelled during 1 hr of collection from *Hoxb8-Cre*^*-/-*^*;Piezo2*^*fl/fl*^ (WT; n=26) and *Hoxb8-Cre*^*+/-*^*;Piezo2*^*fl/fl*^ (KO; n=14) mice (unpaired two-tailed *t*-test: \*\*\**P*=0.0001, t(38)=4.316) (middle right panel). Representative images of dried stools collected during one hour from *Hoxb8-Cre*^*-/-*^*;Piezo2*^*fl/fl*^ (WT) and *Hoxb8-Cre*^*+/-*^*;Piezo2*^*fl/fl*^ (KO) mice (right panel). **C**) Illustration of the *Vil1-Cre;Piezo2* targeting coverage in the GI epithelium, cream color designates *Piezo2* deletion in enterochromaffin cells of intestinal epithelia, but not in nodose, DRG and enteric neurons (left panel). Total GI transit time after carmine red gavage into *Vil1-Cre*^*-/-*^*;Piezo2*^*fl/fl*^ (WT; n=12) and *Vil1-Cre*^*+/-*^*;Piezo2*^*fl/fl*^ (KO; n=5) mice (Mann-Whitney test: *P*=0.7411 two-tailed, U=26.5; not statistically significant) (middle left panel). Number of stools expelled during 1 hr of collection from *Vil1-Cre*^*-/-*^*;Piezo2*^*fl/fl*^ (WT; n=15) and *Vil1-Cre*^*+/-*^*;Piezo2*^*fl/fl*^ (KO; n=9) mice (unpaired two-tailed *t*-test: *P*=0.2591, t(22)=1.158; not statistically significant) (middle right panel). Representative images of dried stools collected during one hour from *Vil1-Cre*^*-/-*^*;Piezo2*^*fl/fl*^ (WT) and *Vil1-Cre*^*+/-*^*;Piezo2*^*fl/fl*^ (KO) mice (right panel). **D**) Illustration of the experimental model to target *Piezo2*-expressing DRG neurons, plum color designates *Piezo2* deletion in DRG neurons, but not in enterochromaffin cells, nodose and enteric neurons (left panel). Total GI transit time after gavaging carmine red into *Piezo2*^*fl/fl*^*::Php*.*s-tdTomato* (Control; n=14) and *Piezo2*^*fl/fl*^*::Php*.*s-iCre* (Cre; n=14) mice (Mann-Whitney test: \*\*\**P*=0.0001 two-tailed, U=20) (middle left panel). Number of stools expelled during 1 hr of collection from *Piezo2*^*fl/fl*^*::Php*.*s-tdTomato* (Control; n=7) and *Piezo2*^*fl/fl*^*::Php*.*s-iCre* (Cre; n=11) mice (Mann-Whitney test: \**P*=0.0208 two-tailed, U=13.5) (middle right panel). Representative images of dried stools collected during one hour from *Piezo2*^*fl/fl*^*::Php*.*s-tdTomato* (Control) and *Piezo2*^*fl/fl*^*::Php*.*s-iCre* (Cre) mice (right panel).

Next, to investigate the concurrent contribution of DRG neurons and gut epithelial cells in GI transit, we used the *Hoxb8*^*Cre*^ driver, which spares nodose ganglia and expresses the Cre recombinase in a gradient pattern targeting cells below the mid-thoracic region ^42^. We validated this driver by crossing it with an *H2b-mCherry* reporter line, which drives nuclear-localized mCherry in Cre expressing cells; to evaluate the recombination efficiency within the ENS, we used whole-mount preparations of mucosal-free intestinal tissues. We confirmed the gradient expression pattern in gut muscle, however nuclei from enteric neurons lacked mCherry expression along the GI tract (Figure S2B), thus *Hoxb8*^*Cre*^ is unable to target enteric neurons. When we assessed the GI function in *Hoxb8*^*Cre+/-*^*;Piezo2*^*fl/fl*^ (*Piezo2*^*Hoxb8*^) mice, we observed accelerated GI transit, increased defecation frequency, increased water content and reduced dry-stool weight in comparison to the wild-type (*Piezo2*^*WT*^) littermates (Figure 3B, Figure S2-E), phenocopying the *Piezo2*^*SNS*^ model. Additionally, the shorter transit time was observed in *Piezo2*^*Hoxb8*^ mice using a videorecorder to detect the colored fecal pellets instead of an experimenter to avoid stress on mice ^43^ (Figure S2-E). These results suggest that Piezo2-expressing intestinal epithelial cells or spinal afferents, rather than enteric or nodose neurons, are responsible for the accelerated GI transit phenotype.

Enterochromaffin cells are a subtype of enteroendocrine cells that have been associated with gut motility ^44,45^. Additionally, Piezo2 is expressed in enterochromaffin cells from the small intestine^39,46^ and colon^40,44,47^, and its deletion was shown to prolong GI transit time in fasted mice ^44,45^. To test whether enterochromaffin cell mechanosensitivity contributes to gut transit in presence of luminal contents, we used the intestinal epithelial *Villin*^*Cre*^ (*Vil1*^*Cre*^) driver to remove Piezo2 from enterochromaffin cells. We observed a similar GI transit time in *Vil1*^*Cre+/-*^*;Piezo2*^*fl/fl*^ (*Piezo2*^*Vil1*^) compared to the wild-type littermate controls (*Vil1*^*Cre-/-*^*;Piezo2*^*fl/fl*^, *Piezo2*^*WT*^) (Figure 3C). Consistently, defecation frequency, water content and dry-stool weight from *Piezo2*^*Vil1*^ mice were all similar to the *Piezo2*^*WT*^ controls (Figure S2-F). These findings suggest that Piezo2 deficiency in enterochromaffin cells is not by itself required for regulating luminal-content transit in *vivo*. Previous studies suggested that Piezo2-defficiency in enterochromaffin cells causes a slight GI transit delay ^44,45^. The difference between these studies might be due to variations in nutrients and microbiota across laboratories. Importantly, as shown above, the accelerated gut transit when Piezo2 is ablated from DRGs and enterochromaffin cells via the *Hoxb8*^*Cre*^ driver, is robust between institutions (Scripps and Mayo Clinic) suggesting a dominant role of DRGs in gut motility.

To determine whether Piezo2 -expressing DRG neurons are responsible for the accelerated transit phenotype, we intrathecally injected peripheral neuron-selective *Php*.*s* viral particles ^48^ carrying a Cre recombinase construct or a fluorescent protein as a control into adult *Piezo2*^*fl/fl*^*/Ai9*^*fl/+*^ mice in between lumbar level 5-6 (Figure 3D, Figure S2-C). This viral strategy was necessary because no existing driver lines target just DRG neurons while sparing nodose and enteric ganglia. Mice with ablated Piezo2 from DRG neurons (*Piezo2*^*DRG*^) presented a profound decrease in the GI transit time in comparison to the wild-type littermate controls (*Piezo2*^*control*^) (Figure 3D). Consistently, the defecation frequency was increased (Figure 3D, rightmost panels). Thus, loss of Piezo2 in DRG neurons is sufficient to drive accelerated GI transit. Notably, as this viral strategy allowed us to induce the phenotype in adult mice, we can exclude the possibility that the accelerated GI transit is consequence of a developmental deficit. These findings indicate that Piezo2 in DRGs is crucial for the maintenance of gut transit homeostasis.

### Neuronal Piezo2 mediates gastric emptying, intestinal transit and colonic transit in mice

Our previous GI transit experiments and others provide information on the time required for intestinal contents to travel from the stomach to the evacuation point ^43,49–51^, but lack details about the transit throughout the intermediate regions of the gut. To investigate whether Piezo2 -expressing somatosensory neurons modulate motility along the entire GI tract or in discrete regions, we functionally evaluated gastric emptying, intestinal transit, and colonic transit. We returned to the *SNS*^*Cre*^*;Piezo2* mouse for these experiments to consistently and uniformly access the majority of the Piezo2-expressing DRG neurons. To probe the function of Piezo2 in gastric emptying, we gavaged *Piezo2*^*SNS*^ and wild-type littermates with a non-absorbable, near-infrared fluorescent dye (GastroSense-750) (Figure 4A). Mice were euthanized at different time points after gavage and the GI tract was harvested and imaged using the IVIS-Lumina S5 system to determine where dye had accumulated. To measure gastric emptying, the fluorescence intensity from the stomach was compared to the rest of the small and large intestines and expressed as percentage of the total signal. We consistently observed faster gastric emptying in *Piezo2*^*SNS*^ mice at 30 min and 45 min after the gavage in comparison to the *Piezo2*^*WT*^ controls (Figure 4B). This indicates that Piezo2 in sensory neurons regulates the rate of stomach emptying.

**Figure 4.**
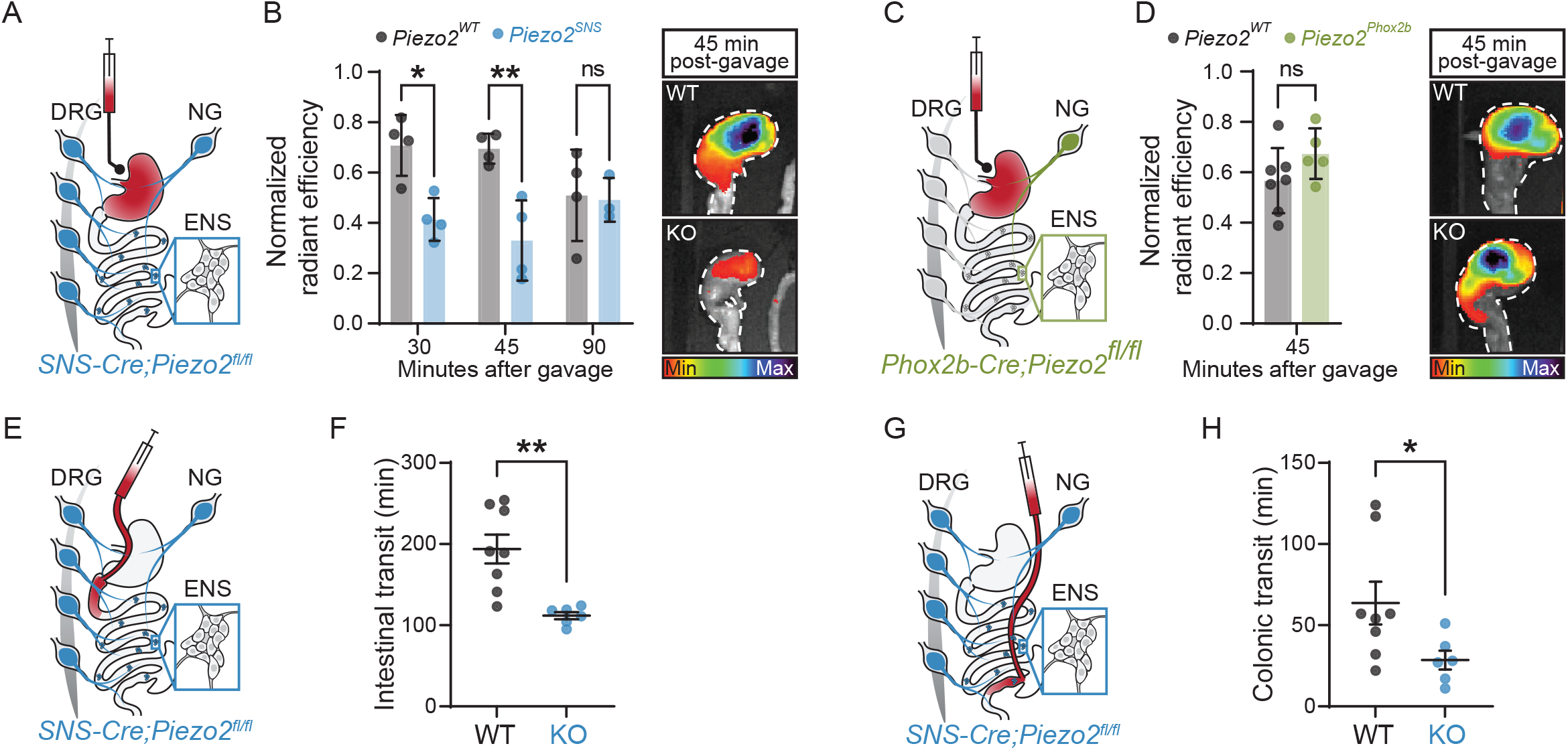
Neuronal Piezo2 mediates gastric emptying, intestinal and colonic transit in mice. **A**) Illustration of the strategy to test gastric emptying in *Piezo2*^*SNS*^ mice. **B**) Quantification of the percentage of gastric emptying observed after gavaging the far-red dye GastroSense-750 at different time points in *SNS-Cre*^*-/-*^*;Piezo2*^*fl/fl*^ (*Piezo2*^*SNS*^; n=3-4 mice per time point) and *SNS-Cre*^*+/-*^*;Piezo2*^*fl/fl*^ (*Piezo2*^*WT*^; n=4 per time point) mice (two-way ANOVA: \*\*\**P*_genotype_=0.0005, *F*(1,17)=18.40; Sidak’s *P*_adjusted_: \**P*_30 min_=0.0122; \*\**P*_45 min_=0.0022; *P*_90 min_=0.9970) (left panel). Representative images of dye emptied from stomach 45 min after gavage *SNS-Cre*^*-/-*^*;Piezo2*^*fl/fl*^ (WT) and *SNS-Cre*^*+/-*^*;Piezo2*^*fl/fl*^ (KO) mice. The stomach is delineated by a white dashed line and pseudocolor scale indicates the dye intensity (right panel). **C**) Illustration of the strategy to test gastric emptying in *Piezo2*^*Phox2b*^ mice. **D**) Quantification of the percentage of gastric emptying observed 45 min after GastroSense-750 gavage in *Phox2b -Cre*^*-/-*^*;Piezo2*^*fl/fl*^ (*Piezo2*^*WT*^; n=7) and *Phox2b -Cre*^*+/-*^*;Piezo2*^*fl/fl*^ (*Piezo2*^*Phox2b*^; n=5) mice (Mann-Whitney test: *P*=0.1061 two-tailed, U=7) (left panel). Representative images of dye emptied from stomach 45 min after gavage *Phox2b-Cre*^*-/-*^*;Piezo2*^*fl/fl*^ (WT) and *Phox2b-Cre*^*+/-*^*;Piezo2*^*fl/fl*^ (KO) mice. Stomach delineated by a white dashed line and pseudocolor scale indicates the dye intensity (right panel). **E**) Schematic of the duodenal infusion in *Piezo2*^*SNS*^ mice through an implanted catheter. **F**) Quantification of intestinal transit time measured after infusing carmine red into the duodenum of *SNS-Cre*^*-/-*^*;Piezo2*^*fl/fl*^ (WT; n=7) and *SNS-Cre*^*+/-*^*;Piezo2*^*fl/fl*^ (KO; n=5) (Mann-Whitney test: \*\**P*=0.0051 two-tailed, U=1). **G**) Schematic of the colonic infusion in *Piezo2*^*SNS*^ mice through a cecum catheter implant. **H**) Quantification of colonic transit time measured after infusing carmine red into the cecum of *SNS-Cre*^*-/-*^*;Piezo2*^*fl/fl*^ (WT; n=8) and *SNS-Cre*^*+/-*^*;Piezo2*^*fl/fl*^ (KO; n=6) (Mann-Whitney test: \**P*=0.0451 two-tailed, U=6).

We previously found that Piezo2 deletion from nodose neurons had no effect on overall GI transit (Figure 3A). Nonetheless, given the importance of vagal innervation in stomach function, we tested the contribution of Piezo2-expressing nodose neurons in gastric emptying. We gavaged *Piezo2*^*Phox2b*^ and wild-type littermates with GastroSense-750 and imaged gut tissues 45 min after gavage (Figure 4C). Consistent with our previous results, we observed no difference in gastric emptying between *Piezo2*^*Phox2b*^ and *Piezo2*^*WT*^ controls (Figure 4D). These results indicate that lack of Piezo2 from nodose neurons is insufficient to accelerate stomach emptying.

Next, we tested whether the small intestine contributes to the accelerated transit observed in *Piezo2*^*SNS*^ mice. We implanted catheters into the duodenum to directly infuse dyes and to quantify the intestinal transit when the stomach is bypassed (Figure 4E). We first infused carmine red through the intestinal catheter and recorded the time until the first colored fecal pellet appeared. We observed a significant decrease in intestinal transit time in *Piezo2*^*SNS*^ compared to wild-type littermate mice (Figure 4F).

These findings reveal that removing Piezo2 from sensory neurons accelerates small intestine transit, suggesting that Piezo2 neurons may be able to modulate small intestine transit independently of stomach emptying activity.

Finally, we directly examined colonic transit by implanting catheters into the cecum to infuse dyes into the proximal colon and circumvent the influence of stomach and small intestine (Figure 4G). When we infused carmine red through the cecal catheter and quantified the time until the first colored fecal pellet appeared, we observed a small but significant decrease in colonic transit time in *Piezo2*^*SNS*^ mice compared to wild-type littermates (Figure 4H). These data show that Piezo2-deficiency in sensory neurons affects the transit of gastric and intestinal contents, indicating that Piezo2-sensory neurons modulate propulsive motility in the stomach, small intestine, and colon in the presence of luminal contents.

### Piezo2-expressing somatosensory neurons innervate the gastrointestinal tract

Next, we examined whether Piezo2-expressing DRG neurons directly project into the GI tract, their morphological endings, and the innervated layer (namely, muscle or mucosa). For this, AAV9 particles encoding a Cre-dependent GFP reporter (*AAV9-flex-GFP*) were injected intrathecally into *Piezo2-ires-Cre* mice (*Piezo2-ires-Cre::AAV9-flex-GFP, Piezo2*^*GFP*^) (Figure 5A). This approach enabled us to specifically visualize Piezo2-DRG endings within the GI tract while sparing vagal and enteric innervation. We mapped and quantified the nerve terminals through image analysis of whole-mount preparations (Figure 5B-C). Interestingly, whole-mount visualization of *Piezo2*^*GFP*^ stomach primarily revealed intraganglionic varicose endings (IGVEs) (Figure 5D). We found no intramuscular arrays or mucosal endings along the GI tract from *Piezo2*^*GFP*^ mice. Although no function has yet been assigned, the IGVE innervation pattern matched previous descriptions of spinal afferents detected in stomach and colon ^52,53^. We observed Piezo2 terminals innervating the small intestine and detected IGVEs and single axons traversing large distances. Further down the GI tract, the colon presented the highest innervation density and the most abundant IGVE network (Figure 5B, C). These findings are consistent with previous studies indicating that spinal innervation is denser towards the large intestine ^28,36^; however, it is important to note that we intrathecally injected between Lumbar levels 5-6, resulting in a gradient pattern of infection with the highest efficiency close to the injection area ^54^ (Figure S3). Therefore, the observed innervation pattern could be additionally explained by our technical approach. Our data revealed that Piezo2 sensory endings from DRG origin innervate the stomach, small intestine, and colon with a predominant morphology of intraganglionic varicose endings.

**Figure 5.**
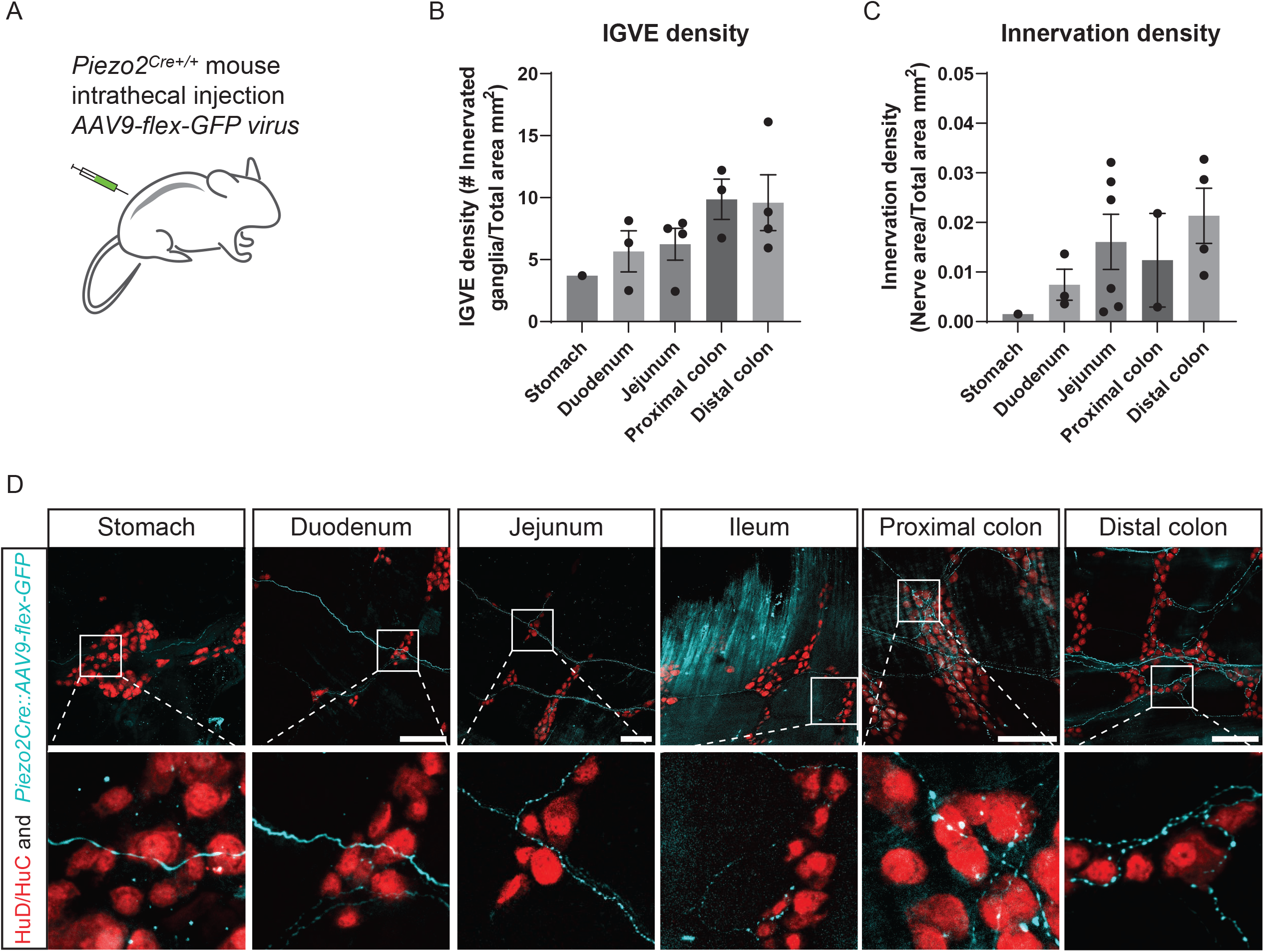
Piezo2 dorsal-root-ganglion neurons innervate the gastrointestinal tract. **A**) Illustration of the strategy to assess DRG neuronal innervation by intrathecally injecting *AAV9-flex-GFP* particles into *Piezo2*^*Cre+/+*^ mice. **B**) Quantification of the IGVE density, defined as the number of enteric ganglia innervated by IGVE in the total area across the whole GI tract.**C**) Quantification of total innervation density, defined as innervated nerve area by the total area across the GI tract.**D**) Representative images of stomach, small intestine, and colon. The enteric neuron nuclei were labeled with HuD/HuC antibody and represented in red. Piezo2-positive nerve endings are shown in cyan. Arrowheads represent the varicosities from the IGVEs.

### Piezo2-expressing DRG neurons detect colon distention

In humans, stool expulsion has been associated with high amplitude propagating contractions that span the entire colon ^55–57^, yet stool evacuation can similarly occur in the absence of this activity by voluntary contracting the abdominal wall and recruiting pelvic floor muscles ^55^. Furthermore, due to the arrival of fecal content, the rectum expands prior to defecation. Nonetheless, *PIEZO2*-deficient individuals perceive the act of evacuation differently because they lack bowel sensation. However, it is unclear whether difficulties in detecting rectal distention affect the overall defecation process. To examine the mouse response to rectum distention, we introduced glass beads into *Piezo2*^*SNS*^ and *Piezo2*^*WT*^ mice and quantify the expulsion time (Figure 6A). It is worth noting that the colonic contents of *Piezo2*^*SNS*^ and *Piezo2*^*WT*^ mice differ in size and water content (Figure S4-A, Figure 2E). The mean diameter of fresh *Piezo2*^*SNS*^ stools is 2.17 mm (± 0.34), which is significantly smaller than the stools from the *Piezo2*^*WT*^ littermates: 2.90 mm (± 0.41) (Figure S4-A). Given these differences, we tested a range of bead sizes. We did not observe any significant difference between *Piezo2*^*SNS*^ and *Piezo2*^*WT*^ littermate mice when 1- and 2-mm beads were used (Figure 6B-C). However, when we used larger 3-mm beads, *Piezo2*^*SNS*^ mice presented a small but significant increase in bead-expulsion time in comparison to the *Piezo2*^*WT*^ littermates (Figure 6D). To confirm the effect of Piezo2 deficiency on rectum motility, we reasoned that an even larger bead (4 mm) would cause a more pronounced motility delay in *Piezo2*^*SNS*^ mice and additionally mimic impacted stools presented in humans who experience constipation. Remarkably, when we tested 4-mm beads, we saw a stark delay in the bead expulsion time in the *Piezo2*^*SNS*^ mice in comparison to the *Piezo2*^*WT*^ controls (Figure 6E). This finding suggests that the lack of Piezo2 impairs the detection of distension, which delays the initiation of mechanically induced peristalsis in rectum. Furthermore, as *Piezo2*^*SNS*^ and *Piezo2*^*WT*^ mice have different stool dimensions, it is possible that Piezo2 neurons have an additional role in regulating stool shape and size.

**Figure 6.**
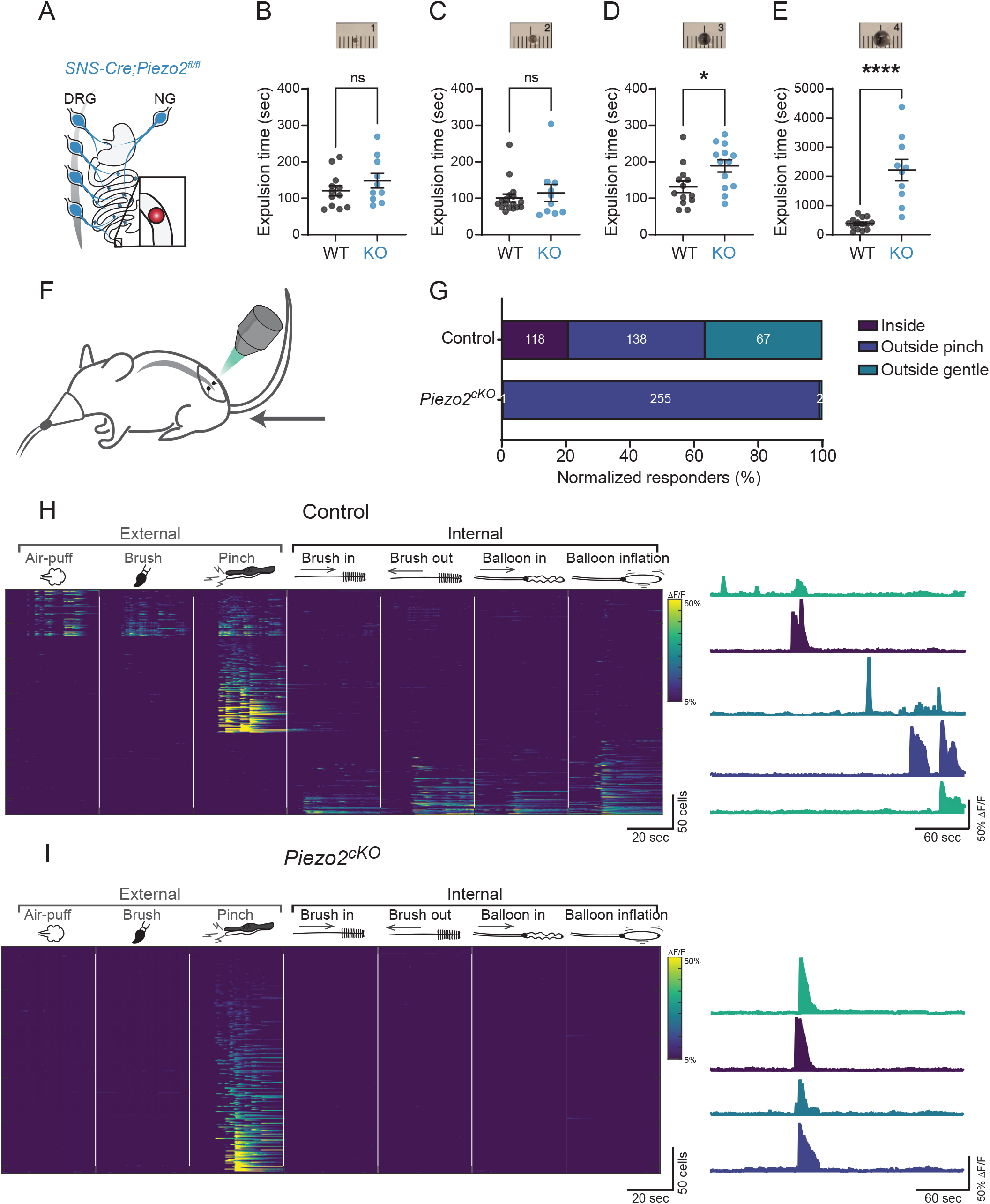
Piezo2 -expressing DRG neurons detect colon distention. **A**) Illustration of the Cre line used for the glass bead expulsion test. **B-D**) Quantification of colon motility test in *SNS-Cre*^*-/-*^*;Piezo2*^*fl/fl*^ (WT) and *SNS-Cre*^*+/-*^*;Piezo2*^*fl/fl*^ (KO) using different sizes of glass beads. A picture a representative bead is shown above the quantification.**B**) 1 mm bead (unpaired two-tailed *t*-test: *P*=0.2592, t(20)=1.161; ns, not statistically significant).**C**) 2 mm bead (Mann-Whitney test: *P*=0.9900 two-tailed, U=84.5; not statistically significant).**D**) 3 mm bead (unpaired two-tailed *t*-test: \**P*=0.0196, t(24)=2.500). **E**) 4 mm bead (unpaired two-tailed *t*-test: \*\*\*\**P*<0.0001, t(22)=5.910). **F**) Illustration of *in vivo* calcium imaging recording in anesthetized mice. **G**) Quantification of the responses obtained from Control (*Hoxb8*^*Cre*^*;GCaMP6f*^*+/+*^, n=4) and *Piezo2*^*cKO*^ (*Hoxb8*^*Cre*^*;Piezo2*^*fl/fl*^*;GCaMP6f*^*+/+*^, n=4). The insets represent the numbers of recorded cells per condition. **H**) Heatmap showing calcium responses recorded from Control (*Hoxb8*^*Cre*^*;GCaMP6f*^*+/+*^) DRG neurons (n=323 cells ; N=4 mice). External and internal stimulations are shown. Representative traces are shown on the right. **I**) Heatmap showing Calcium responses recorded from *Piezo2*^*cKO*^ (*Hoxb8*^*Cre*^*;Piezo2*^*fl/fl*^*;GCaMP6f*^*+/+*^) DRG neurons (n=258 cells ; N=4 mice). External and internal stimulations are shown. Representative traces are shown on the right.

So far, our findings indicate that Piezo2-positive DRG fibers are present throughout the GI tract, and that motility is affected in Piezo2-deficient mice in all investigated gut regions. Next, we asked whether Piezo2 is directly required to sense mechanical stimulation within the gut. For this purpose, we adopted a novel colon preparation where we introduce a soft brush and inflate a balloon into the colon of anesthetized mice, while simultaneously recording DRG neuron activity using the calcium-sensitive indicator GCamp6f (Figure 6F). For these experiments, we used *Hoxb8*^*Cre+/+*^*;Piezo2*^*fl/fl*^*;GCaMP6f*^*+/+*^ (*Piezo2*^*cKO*^) mice and as control *Hoxb8*^*Cre*^*;GCaMP6f*^*+/+*^ (*Piezo2*^*WT*^). This approach enabled us to monitor the calcium signal from sacral DRG neurons in *Piezo2*^*cKO*^ and wild-type littermates. As a control response, we externally stimulated the perineal region with a puff of air, a gentle brush stroke, and a pinch. As internal stimulation, we utilized a soft brush movement and inflated a balloon inside the colon. We hypothesized that DRG neurons expressing Piezo2 detect colon stretch to allow calcium influx. *Piezo2*^*WT*^ mice exhibited rapid and robust responses in sacral level 1 (S1) neurons after the external stimulation with an air puff, a gentle stroke and a noxious pinch in the anal area (Figure 6G-H). We additionally observed calcium responses when introducing and removing a soft brush into the colon, and after inflating a colonic balloon in *Piezo2*^*WT*^ mice (Figure 6H). Furthermore, all the calcium responses were segregated in internal and external stimuli supporting that different DRG neurons innervate perineal skin and colon. Consistent with previous findings ^58^, responses to gentle stimuli (air puff and brush stroke) were markedly attenuated in somatosensory neurons from *Piezo2*^*cKO*^ mice (Figure 6G, I), corroborating the role of Piezo2 in the sense of touch. Strikingly, all responses to colonic stimuli (brush insertion and extraction, and balloon inflation) were abolished in DRG neurons from *Piezo2*^*cKO*^ mice, and only the response to painful pinch remained (Figure 6G, I). This indicates that Piezo2 from DRG neurons is a key sensor of colon stretch.

## Discussion

The importance of gut motility and its control has been recognized since the 18th century ^7^. The GI tract is extensively innervated by the enteric nervous system ^18,19,59^, vagal afferents ^8,9^, and somatosensory neurons of the thoracic, lumbar, and sacral DRG ^28,60^. Here we demonstrate that ingested contents provide mechanical feedback through activation of Piezo2 to dramatically slow the gut transit. Remarkably, using an array of conditional knockout mice, we uncovered that this food-dependent brake relies exclusively and unexpectedly on DRG mechanosensory input.

Whereas gut transit plays a major role in efficient digestion and nutrient absorption, defecation is another critical function of the lower GI tract that is known to be independently controlled ^61^. Notably, Piezo2 knockout mice exhibited a delayed evacuation in a bead-expulsion assay and exhibited diarrhea-like behavior, possibly due to a failure to resorb water caused by the reduced transit times.

Interestingly, human subjects with PIEZO2 deficiency also exhibit frequent GI dysregulation that ranges from constipation to diarrhea, consistent with the diverse roles of Piezo2 in controlling gut motility and defecation in mice.

Our data has therapeutic implications for a range of GI disorders. We anticipate that inhibition and activation of PIEZO2 could enhance or slow gut transit, respectively. Furthermore, using *in vivo* functional imaging, we found that Piezo2 is essential for all types of mechanosensation by DRG neurons innervating the colon in male and female mice. It is notable that stimulation using balloon inflation produces forces well into the noxious range ^62–64^. Intriguingly, conditional deletion of Piezo2 in neurons expressing *Scn10a* produced similar phenotypes to broadly knocking out this mechanoreceptor for all DRG neurons, suggesting a potential role of Piezo2 in gut mechano-nociception.

Taken together, our data provide a molecular and cellular explanation for how gut contents trigger mechanosensory-DRG neurons to control transit through the GI system. Whether Piezo2 in sensory endings detect the luminal contents passing through the gut or the constant gut contractions triggered by luminal contents is currently unknown. Furthermore, a key unanswered question remains as to how activation of DRG neurons decrease gut motility? A clue will be uncovered by determining the anatomical organization and central projections of the specific classes of Piezo2-expressing DRG neurons targeting the gut. Our study focused on the role of Piezo2 in the somatosensory system.

Future studies should reveal the role of this mechanosensitive ion channel in the vagal and enteric neurons. Lastly, it has been shown that the sensitivity of gut innervating mechanosensory neurons can be significantly sensitized by inflammation common to a range of GI disorders ^65–67^. Most notably Inflammatory Bowel Disease (IBD), that can be extremely painful, causes diarrhea or constipation, and yet we lack effective treatment. Determining how PIEZO2 function is altered during gastrointestinal disease will be particularly important.

## Supplementary figures

**Figure Supplementary 1.**
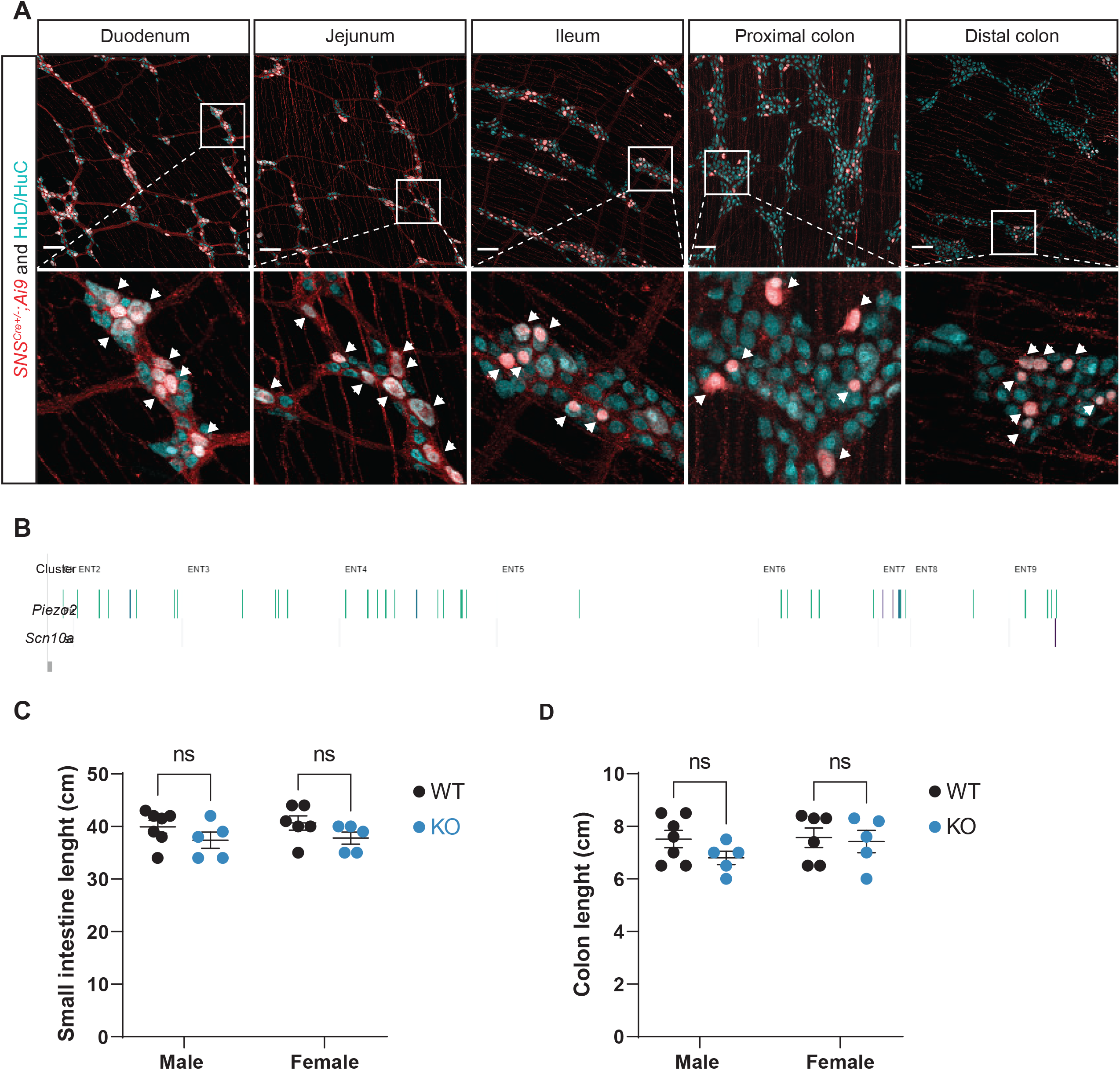
ENS validation of *SNS-Cre*^*+/-*^*;Ai9*^*fl/fl*^ mice along the GI tract. **A**) Representative images from whole-mount preparations of small and large intestine. Enteric neuron nuclei are labeled with HuD/HuC and represented in cyan. *SNS* positive fibers are represented in red. **B**) Comparison of *Piezo2* and *Scn10a* transcript from enteric neurons, taken from ^38^. **C**) Comparison of small intestine length from *SNS-Cre*^*-/-*^*;Piezo2*^*fl/fl*^ (WT; n=12) and *SNS-Cre*^*+/-*^ *;Piezo2*^*fl/fl*^ (KO; n=10) mice. **D**) Comparison of colon length from *SNS-Cre*^*-/-*^*;Piezo2*^*fl/fl*^ (WT; n=12) and *SNS-Cre*^*+/-*^*;Piezo2*^*fl/fl*^ (KO; n=10) mice.

**Figure Supplementary 2.**
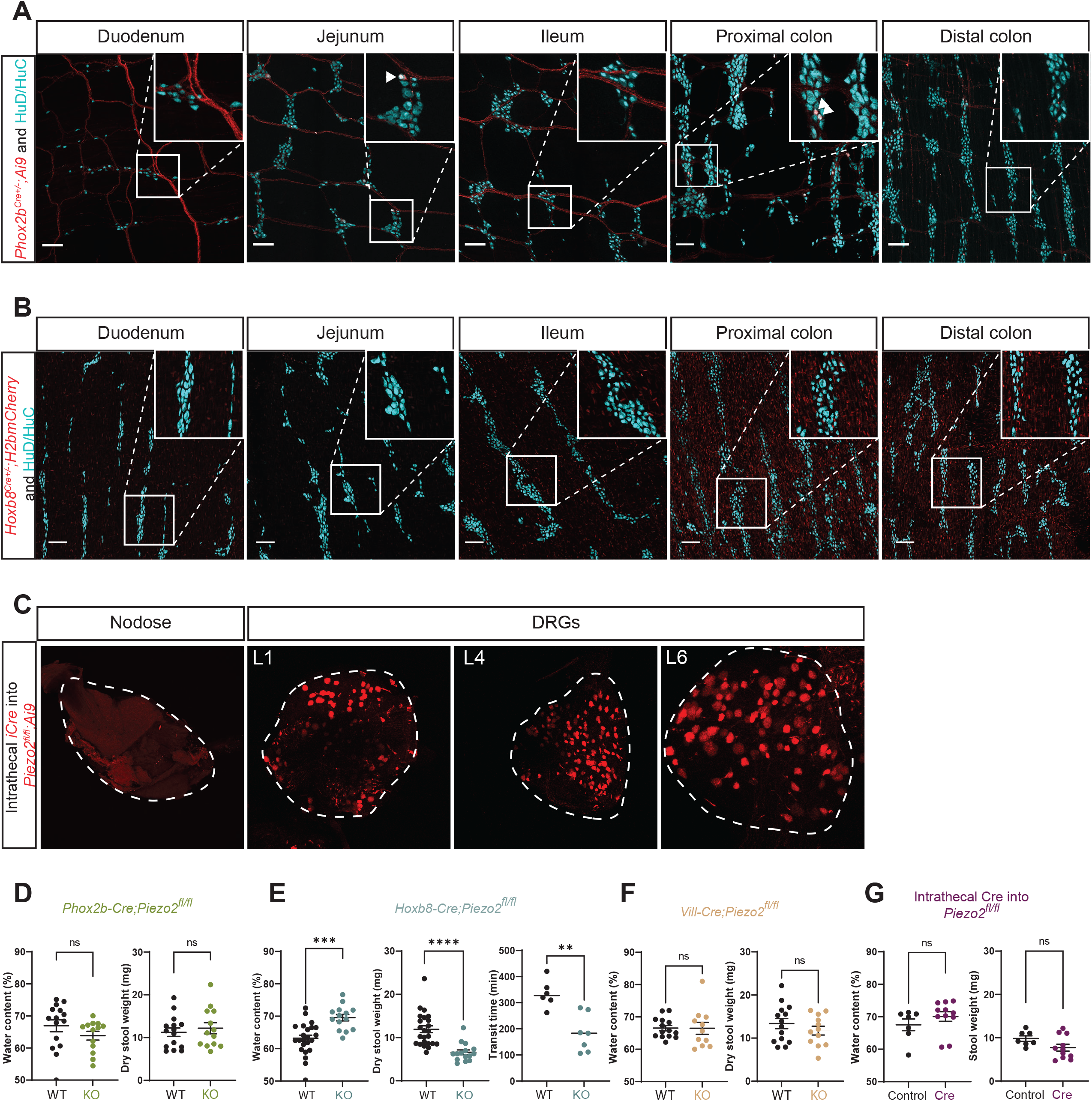
Validation of *Phox2b-Cre*^*+/-*^*;Ai9*^*fl/fl*^, *Hoxb8-Cre*^*+/-*^*;H2bmCherry* and *Piezo2*^*fl/fl*^ intrathecally injected with *Php*.*s-iCre*. **A**) Representative images of whole-mount preparation of small and large intestine. Enteric neuron nuclei are labeled with HuD/HuC and represented in cyan. *Phox2b* positive fibers are represented in red. **B**) Representative images of whole-mount preparation of small and large intestine. Enteric neuron nuclei are labeled with HuD/HuC and represented in cyan. *Hoxb8* positive signal is represented in red. **C**) Representative images of nodose and DRGs after intrathecal injection with *Php*.*s-iCre*. **D**) Quantification of stool water content and dry stool weight from *Phox2b-Cre*^*-/-*^*;Piezo2*^*fl/fl*^ (WT; n=15) and *Phox2b-Cre*^*+/-*^*;Piezo2*^*fl/fl*^ (KO; n=13) mice. **E**) Quantification of stool water content from *Hoxb8-Cre*^*-/-*^*;Piezo2*^*fl/fl*^ (WT; n=25) and *Hoxb8-Cre*^*+/-*^ *;Piezo2*^*fl/fl*^ (KO; n=14) mice (unpaired two-tailed *t*-test: \*\*\**P*=0.0001, t(37)=4.253) (left panel). Quantification of dry stool weight from *Hoxb8-Cre*^*-/-*^*;Piezo2*^*fl/fl*^ (WT; n=25) and *Hoxb8-Cre*^*+/-*^*;Piezo2*^*fl/fl*^ (KO; n=15) mice (unpaired two-tailed *t*-test: \*\*\*\**P*<0.0001, t(38)=4.857) (center panel). Quantification of GI transit time from *Hoxb8-Cre*^*-/-*^*;Piezo2*^*fl/fl*^ (WT; n=6) and *Hoxb8-Cre*^*+/-*^*;Piezo2*^*fl/fl*^ (KO; n=7) mice (Mann-Whitney test: \*\**P*=0.0047, two-tailed, U=2) (right panel). **F**) Quantification of stool water content and dry stool weight from *Vil1-Cre*^*-/-*^*;Piezo2*^*fl/fl*^ (WT; n=14) and *Phox2b-Cre*^*+/-*^*;Piezo2*^*fl/fl*^ (KO; n=11) mice. **H**) Quantification of stool water content and dry stool weight from Control (n=7) and *iCre* (n=11) mice.

**Figure Supplementary 3.**
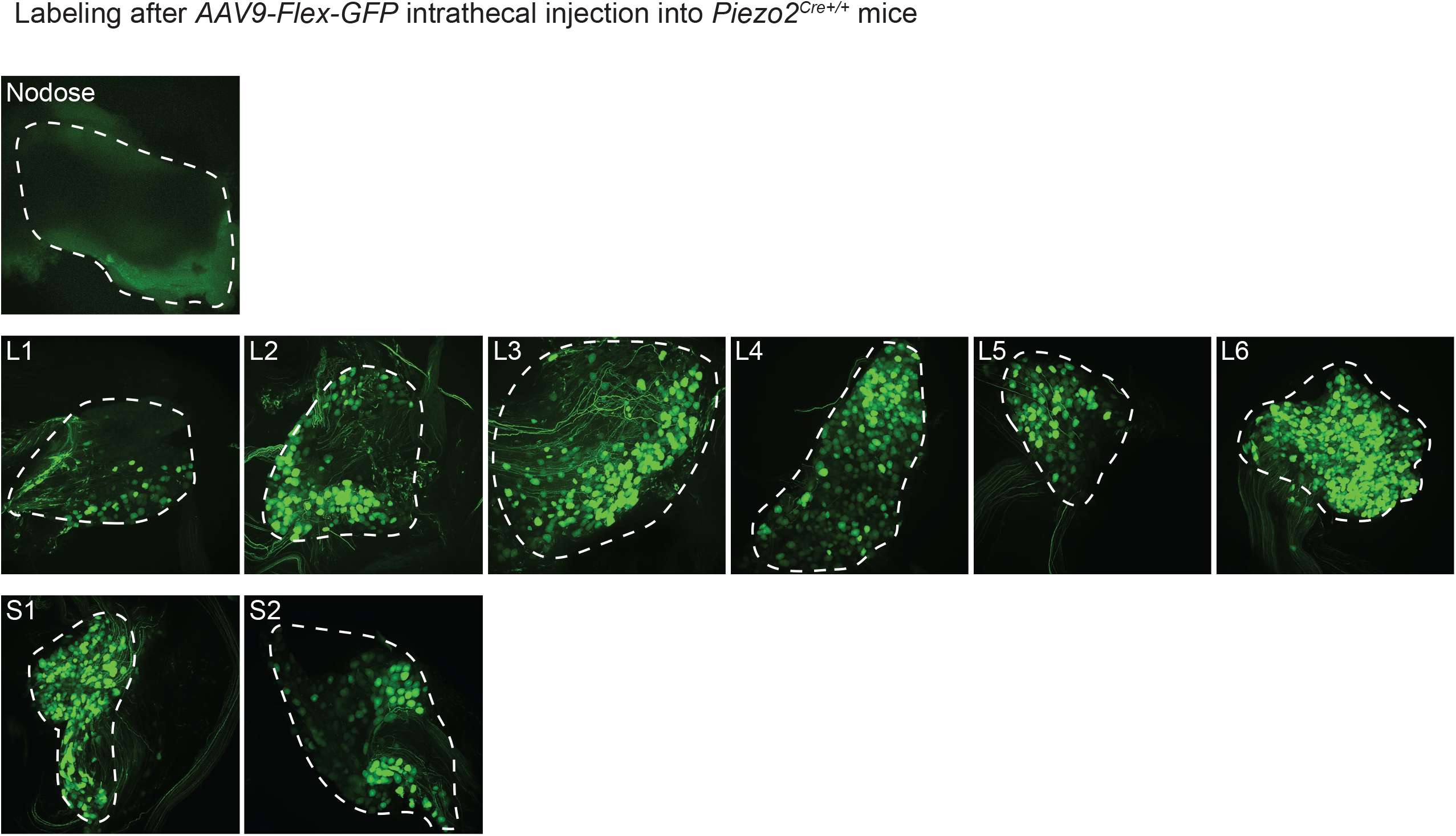
DRG validation after intrathecal injection of *AAV9-flex-GFP* particles into *Piezo2-Cre*^*+/+*^ mice. Representative images of whole-mount preparation of nodose and DRGs 4 weeks after intrathecal injection of *AAV9-flex-GFP* particles into *Piezo2-Cre*^*+/+*^ mice.

**Figure Supplementary 4.**
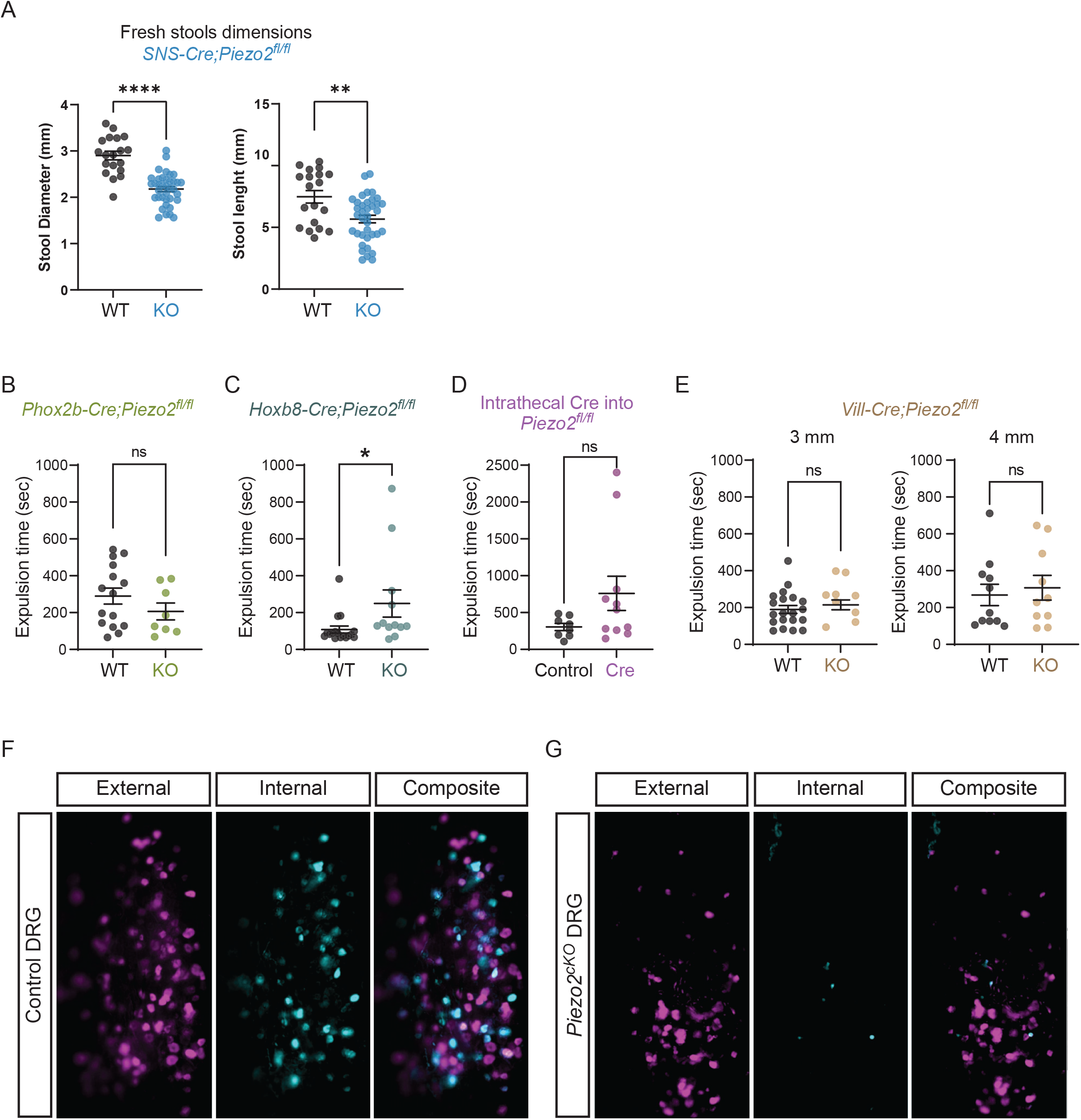
DRG validation after intrathecal injection of *AAV9-flex-GFP* particles into *Piezo2-Cre*^*+/+*^ mice. **A**) Left, quantification of stool diameter collected from *SNS-Cre*^*-/-*^*;Piezo2*^*fl/fl*^ (WT; n=19 ; N=4 mice) and *SNS-Cre*^*+/-*^*;Piezo2*^*fl/fl*^ (KO; n= 39; N=3 mice) mice (unpaired two-tailed *t*-test: \*\*\*\**P*<0.0001, t(54)=7.037). Right, quantification of stool length collected from *SNS-Cre*^*-/-*^*;Piezo2*^*fl/fl*^ (WT; n=19 ; N=4 mice) and *SNS-Cre*^*+/-*^*;Piezo2*^*fl/fl*^ (KO; = 39; N=3 mice) mice (unpaired two-tailed *t*-test: \*\**P*=0.0021, t(54)=3.236). **B**) Colon motility test in *Phox2b-Cre*^*-/-*^*;Piezo2*^*fl/fl*^ (WT; n=15) and *Phox2b-Cre*^*+/-*^*;Piezo2*^*fl/fl*^ (KO; n=8) using 3 mm bead (unpaired two-tailed *t*-test: *P*=0.2401, t(21)=1.209; ns, not statistically significant).**C**) Colon motility test in *Hoxb8-Cre*^*-/-*^*;Piezo2*^*fl/fl*^ (WT; n=17) and *Hoxb8-Cre*^*+/-*^*;Piezo2*^*fl/fl*^ (KO; n=12) using 3 mm bead (unpaired two-tailed *t*-test: *P*=0.0409, t(27)=2.147; ns, not statistically significant).**D**) Colon motility test in *Piezo2*^*fl/fl*^*::Php*.*s-tdTomato* (Control; n=8) and *Piezo2*^*fl/fl*^*::Php*.*s-iCre* (Cre; n=10) mice (Mann-Whitney test: *P*=0.1288 two-tailed, U=25; ns, not statistically significant). **E**) Colon motility test in *Vil1-Cre*^*-/-*^*;Piezo2*^*fl/fl*^ (WT; n=20) and *Vil1-Cre*^*+/-*^*;Piezo2*^*fl/fl*^ (KO; n=14) using 3 mm bead (unpaired two-tailed *t*-test: *P*=0.4737, t(32)=0.7250; ns, not statistically significant) (left panel). Colon motility test in *Vil1-Cre*^*-/-*^*;Piezo2*^*fl/fl*^ (WT; n=11) and *Vil1-Cre*^*+/-*^*;Piezo2*^*fl/fl*^ (KO; n=9) using 4 mm bead (unpaired two-tailed *t*-test: *P*=0.6622, t(19)=0.4438; ns, not statistically significant) (right panel). **F**) Representative images of standard deviations from all images corresponding to external stimuli and all internal stimuli corresponding to one Control mouse. **G**) Representative images of standard deviations from all images corresponding to external stimuli and all internal stimuli corresponding to one *Piezo2*^*cKO*^ (*Hoxb8*^*Cre*^*;Piezo2*^*fl/fl*^*;GCaMP6f*^*+/+*^) mouse.

## Methods

## Key Resource Table

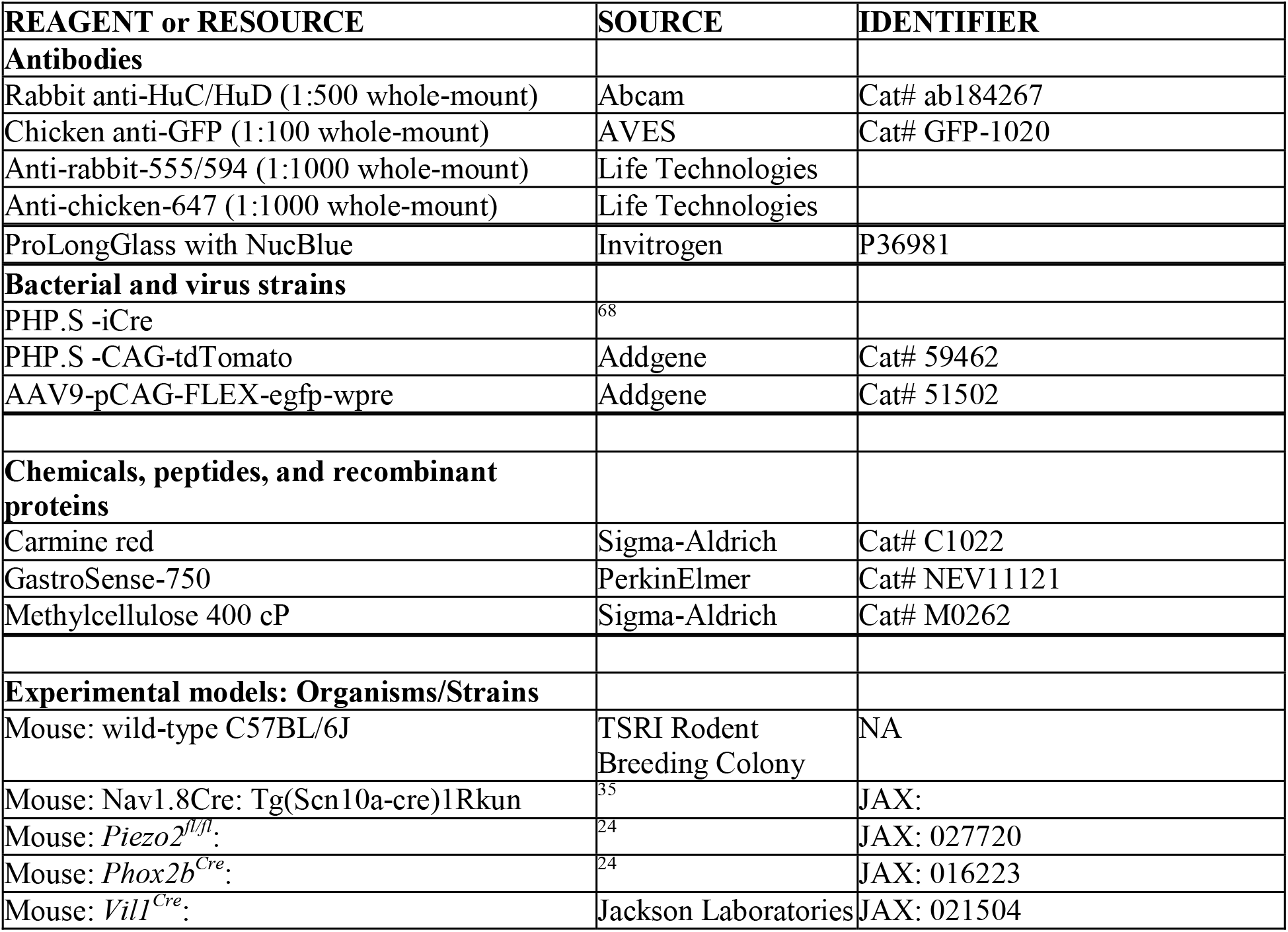
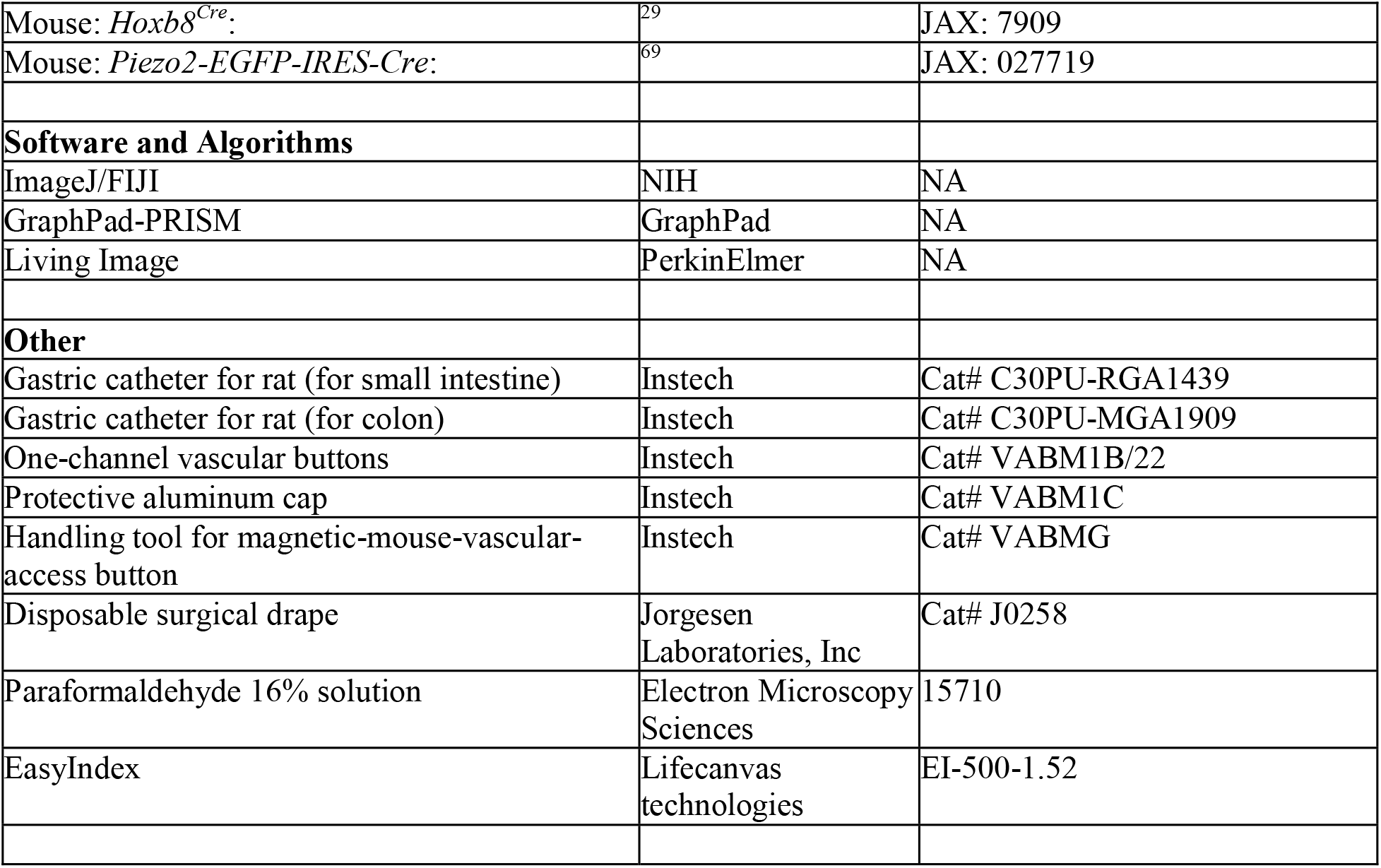

## Lead contact and materials availability

Further information and request for reagents and recourses should be directed to A. Patapoutian or A.T. Chesler.

This study did not generate new unique reagents.

## Experimental model and subject details

Mice were group housed in standard housing under 12–12□hr light–dark cycle and *ad libitum* access to water and standard chow unless noted otherwise. Room temperature was kept at around 22□°C and humidity between 30–80% (not controlled). Mice were kept on pelleted paper bedding and provided with nesting material and a polyvinyl chloride pipe enrichment. Age-matched littermates between 2 and 5 months were used for all *in vivo* experiments. All studies employed a mixture of male and female mice. All the experimental protocols were approved by The Scripps Research Institute Institutional Animal Care and Use Committee and were in accordance with the guidelines from the NIH.

## Clinical assessment

Seven patients with *PIEZO2* loss-of-function mutations were surveyed evaluated at the National Institutes of Health (NIH) under research protocol approved by the Institutional Review Boards of National Institute of Neurological Disorders and Stroke (NINDS, protocol 12-N-0095) between April 2015 and May 2020. Patients were recruited from all over the world and their age ranged between 9 to 42 years at the time of the evaluation. The subject identifier published in the current study corresponds to the same identifier used for the urinary function ^25^. Written informed consent and/or assent (for minor patients) was obtained from each participant in the study. Detailed history, clinical evaluation and testing have been previously described ^25^. PROMIS questionnaires, a clinical tool developed by the National Institute of Health ^30,31^, were used to capture general GI symptoms from the seven days prior to the survey. Parents assisted with information gathering from their children.

## Recombinant viruses

PHP.S-Cre, was obtained from Janelia, PHP.S-TdTomato plasmid was obtained from Addgene (59462). PHP.S particles were produced in-house, titrated by qPCR and aliquoting into 5μl and flash-frozen for long-term storage. AAV9 particles were acquired from Addgene.

## Surgeries

Mice were anaesthetized with isoflurane (4% for induction and 1.5–2% for maintenance) and kept on a heating pad during the procedure. Ophthalmic ointment was applied to the eyes. Skin at the surgical area was shaved, hair removed and sterilized using ethanol and iodine. After surgery, mice were transferred to a warm cage to recover, subcutaneous injection of flunixin was given for 2 days and topical antibiotic ointment was used for post-operative care.

## Intrathecal injections

Mice were injected at 6-7 weeks of age. After pre-surgical care, a 1.5 cm incision was made starting at the level of femur-hip connection extending towards through the midline of the back towards the head. 7 μl of viral particles in PBS with 0.001% F-68 and 0.01% FastGreen were injected into the L5-L6 intervertebral space using a 25μl Hamilton syringe. The skin was closed, and post-surgical care was provided.

For whole-mount analysis, mice were injected with *AAV9-flex-GFP* (1×10^13^ VG per ml, 7 μl) and recovered for a minimum of 4 weeks before perfusion. For GI transit assessment, mice were injected with *PHP*.*S -iCre* or *PHP*.*S -CAG-tdTomato* (1×10^13^ VG per ml, 7 μl) allowed to recover for a minimum of 4 weeks before behavior tests. Consistent with previous studies on the role of Piezo2 in proprioception ^21^, we observed that 8 out of 11 *Piezo2*^*DRG*^ mice lacked proprioception in their hindlimbs. All 11 mice were included in the analysis.

## Intra-intestinal catheter implantation

For this procedure, mice had at least 8 weeks of age. Mice anesthetized with isoflurane, pre-surgical care, and aseptic preparation was taken. An abdominal midline incision through the skin and muscle was performed, extending from the xyphoid process about 1.5 cm caudally. A second 1-cm incision was made between the scapulae for catheter externalization. The skin was separated from the subcutaneous tissue to form a subcutaneous tunnel between the neck and abdomen incisions to facilitate catheter placement. A small puncture hole was made on the left side of the abdominal wall to insert the catheter (Instech, C30PU-RGA1439). The stomach was externalized, and a purse-string stitch was made at the edge of the fundus and corpus on the side of the greater curvature of the stomach using 7-0 non-absorbable Ethilon suture. Then, a puncture was made at the center of the purse-string stitch to insert and advanced the catheter 2.5 cm distal to the pyloric sphincter (intraduodenal catheter). While for the intracecal catheter, a puncture was made on the larger curvature of the cecum to insert and advance the catheter 1 cm, at the edge of the colon and cecal junction. The cecal catheter was secured to the tissue with sterile surgical drape (Jorgesen Laboratories, J0258). The catheter was secured by the purse-string suture at the catheter collar. The abdominal cavity was irrigated with sterile saline and the abdominal wall was closed. The other end of the catheter was attached to a vascular button (Instech, VABM1B/22), sutured to the muscle layer at the interscapular site and the incision was closed. The vascular button was closed with a protective aluminum cap (Instech, VABM1C) to prevent catheter obstruction. Mice were provided with subcutaneous flunixin and moistened chow for 2 days after surgery. Mice were allowed to recover for 7-10 days prior to behavioral experiments.

## Treatments

### Intraduodenal and intracecal infusions

All mice were fed *ad libitum* before experiments and solutions were infused via intraduodenal or intracecal catheters using a handling tool for the vascular button (Instech, VABMG). 100 μl and 50 μl of carmine red was infused through the intraduodenal and intracecal catheter respectively.

### Oral gavage

Mice were gavaged with volumes ranging from 100-300 μl of carmine red or GastrSense-750. All gavages were performed between 8:00-9:00 am. After carmine-red gavage, mice were monitored every 15 min for the presence of the first red fecal pellet.

## Histology

### Whole-mount preparation of GI tissues, nodose and DRGs

Mice were terminally anaesthetized with isoflurane, euthanized by cervical dislocation, and intracardially perfused with ice-cold PBS and ice-cold 4% PFA (Electron Microscopy Perfusion Fixative, 1224SK). Nodose and DRGs were extracted and post-fixed in 4% PFA for 1 hr before being washed with PBS. Nodose and DRGs were mounted onto silicone isolators (Electron Microscopy Sciences, 70345-39) and mounted using EasyIndex (lifecanvas technologies, ei-500-1.52).

Gastrointestinal tissues were extracted, washed with PBS to remove all intestinal contents. Gut tissues were opened and pinned onto syligard-coated dishes. Gut samples were post-fixed in 4% PFA at 4□°C overnight before being washed in PBS. The mucosa was carefully dissected from the muscularis. Tissues were blocked with gentle agitation for 2 hrs at room temperature in (5% normal goat serum, 20% DMSO, 75% PBST (PBS with 0.3% TritonX-100)). Primary antibodies were added to the blocking buffer at appropriate concentrations and incubated for two days at 4□°C. Tissues was washed 3 times in PBST and then incubated in blocking buffer with secondary antibodies overnight at 4□°C. Samples were again washed 3 times in PBST and mounted with ProLongGlass with NucBlue.

## Imaging

### Confocal microscopy

Mounted nodose and DRGs samples were imaged on either a Nikon C2 or Nikon AX scope confocal microscope using a 20x/0.75 LNA objective or a 16x/0.80W respectively. Images were acquired using NIS-Elements.

## Behavioral and physiological assays

### Whole gastrointestinal transit

Mice fed *ad libitum* were gavaged with 300 μl of carmine red (6% carmine red in 0.5% 400 cP methylcellulose) and placed individually into clean cages with access to chow pellets and water.

For experiments comparing GI transit between fasted and fed conditions, mice were fasted 12 hrs prior the gavage with free access to water. Mice were gavaged with 500 μl of carmine red and placed individually into clean cages with access to water but no access to food. Seven days later, same mice were fed *ad libitum*, gavaged with 500 μl of carmine red and placed individually into clean cages with access to chow pellets and water.

### Small and large intestinal transit

Mice fed *ad libitum* were infused with 100 μl or 50 μl of carmine red into intraduodenal or intracecal catheter respectively. Mice were placed individually into clean cages with access to chow pellets and water. After infusions, mice were monitored every 15 min for the presence of the first red fecal pellet.

### Colon motility assay

Mice fed *ad libitum* were anesthetized with isoflurane, then a glass bead was inserted 2 cm into the colon with a gavage cannula. The time for the bead release was recorded, every experiment was performed twice with 7 days in between trials and results were averaged. Only mice that recovered from anesthesia within 60 sec were included in the quantification. For the 4-mm bead experiment, mice weighted at least 24 gr.

### Gastric emptying evaluation

A day prior to the experiment, all hair was removed from the abdominal area. Mice fed *ad libitum* were gavaged with 100 μl of GastroSense-750. Mice were anesthetized with isoflurane 30, 45 and 95 min after gavage. The full GI tract was harvested and immediately imaged using the IVIS-Lumina S5 system. For analysis, Living Image (Perking Elmer) software was used to draw ROIs delineating the stomach and the rest of the GI tract. The radiant efficiency from the stomach was compared to the rest of the small and large intestines and expressed as percentage of the total signal.

### *In vivo* epifluorescence calcium imaging of sacral ganglia

Mice were placed on a mesh floor for 1hr to defecate freely. Scruffing and lower abdomen massage was applied before being anesthetized with isoflurane and transferred to a custom platform. Lower limbs and tail were restrained on this platform and hand warmers were used to maintain body temperature. The dorsal aspect of the sacrum was surgically exposed after partial removal of the gluteus medius and stabilized with a spinal clamp (Narishige STS-A).Using a dental drill, the dorsal root ganglia in the pelvis was exposed by removing a portion of the auricular surface along with the posterior articular process of the 6th Vertebra and the posterior articular process (S2); hemostatic dental sponges (Pfizer Gel Foam) were applied as needed to control bleeding. Following surgery, the animal was transferred to the stage of a custom tilting light microscope (Thorlabs Cerna) equipped with a 4X, 0.28 NA air objective (Olympus). GCaMP6f fluorescence images were acquired with a CMOS camera (PCO Panda 4.2) using a standard green fluorescent protein (GFP) filter cube in 40 second epochs at 5hz. External mechanical stimuli was applied to the animal skin around the anus included a series of pressurized air puffs from a Picospritzer (25psi, for 0.2, 1, 3 and 5 seconds), manual gentle brushing with an acrylic brush and skin pinch with forceps (Students). Internal mechanical stimuli were applied by placing a lubricated gavage tip 4 cm into the rectum and snaking the tip of a dental brush flosser attached to a wire through it. The brush was then pushed out of the gavage tip 2 cm for a total colon depth of 6 cm. Lubricated custom balloons, also built on the backbone of the gavage tip, were placed 6cm into the distal colon of the mouse and inflated until an internal pressure of 100 mmHg, 150 mmHg and 200 mmHg were reached.

Analysis of calcium imaging was performed as previously described ^70^. Regions of interest (ROI) outlining responding cells were drawn in FIJI/ImageJ and relative change of GCaMP6f fluorescence was calculated as percent ΔF/F. Contaminant signal e.g., from out-of-focus tissue and neighboring cells was removed by subtracting the fluorescence of a donut-shaped area surrounding each ROI using a custom MATLAB script. Overlapping ROIs and rare spontaneously active cells were excluded from the analysis. Imaging episodes were concatenated for display as traces or activity heatmaps. Spatial maps of activity were generated by calculating the standard deviation for each pixel over a stimulation episode in FIJI/ImageJ as described previously.

### Quantification of nerve density in stomach, small intestine and colon

Regions of 80□μm □× □80□μm □× □20 □μm (x,y,z) were randomly selected and maximally projected over z using customized ImageJ scripts in the whole stacks of stomach, small intestine and colon from *Piezo2*^*Cre+/+*^ mice that were intrathecally injected with *AAV9-flex-GFP*. Areas containing nerve fibers were automatically segmented using auto thresholding in ImageJ. IGVE were quantified manually as LabeledGanglia/TotalArea. Nerve density was calculated as NerveArea/TotalArea. Only views containing nerve signals were retained for quantification. We quantified 656 views from 39 images from 6 biological replicates.

### Quantification and statistical analysis

Data were expressed as means ± SEMs in figures and text. Normality tests were used, and parametric or non-parametric tests were performed as appropriate. Unpaired two-tailed t tests or Mann-Whitney test were performed. Two-way ANOVA was used to make comparisons across more than two groups using Prism 9.4 (GraphPad). Test, statistics, significance levels, and sample sizes for each experiment are listed on the figure legends.

## Acknowledgements

We thank M. Szczot and Felipe Meira de Faria for their support to develop the *in vivo* colonic preparation. We thank R.Hill, R.Pak and A.Dubin for their feedback on the manuscript. We also thank K. Spencer, the Nikon Center of Excellence Imaging Center, and the Scripps Research Department of Animal Resources for support services. This work was supported by the Howard Hughes Medical Institute, NIH grant R35 NS105067 (AP), and by the intramural program of the NIH, the National Center for Complementary and Integrative Health and National Institute of Neurological Disorders and Stroke (ATC).

## Contributions

M.R.S.V., A.P. and A.T.C. conceived and designed the study. M.R.S.V., R. L., A.K., Y.W., M.L., and H.K. performed experiments and analyzed data. R.M.L. performed and analyzed *in vivo* DRG recordings. H.K. performed and analyzed videorecorded GI transit experiments. M.R.S.V., A.P. and A.T.C. wrote the manuscript. All authors provided input and reviewed the manuscript.

